# A Deep Learning Framework for Quantifying Dynamic Self-Organization in *Myxococcus xanthus*

**DOI:** 10.1101/2025.11.07.686839

**Authors:** Jiangguo Zhang, Eduardo A. Caro, Peiying Chen, Trosporsha Tasnim Khan, Patrick A. Murphy, Lawrence J. Shimkets, Ankit B. Patel, Roy D. Welch, Oleg A. Igoshin

## Abstract

Under starvation, *Myxococcus xanthus* bacteria initiate a multicellular developmental program in which cells move to form fruiting bodies and differentiate into distinct cell types. Many genes affecting this process have been identified, and it is assumed that perturbing genes within the same pathway induces similar changes in the phenotype, although those changes may be subtle or obscured by pleiotropic effects. However, these pathways cannot be systematically mapped, as there are no systematic methods for quantifying phenotype similarity. Here, we applied deep learning techniques to quantify the phenotype patterns and self-organization dynamics of 292 genetically distinct strains. By integrating ResNet and StyleGAN2 to construct a Variational AutoEncoder (VAE) and utilizing a Siamese network for phenotypic similarity metrics, we efficiently encoded high-resolution microscopy images into 13-dimensional feature vectors, capturing phenotypic variability in aggregation patterns across time and strains. Human evaluation confirmed that our model’s reconstructions were visually indistinguishable from real images and closely aligned with input phenotypes. Importantly, the feature space is interpretable: individual dimensions correlate with biological features such as aggregate number and size, and extrapolation along these dimensions produces predictable morphological changes. Remarkably, our model revealed that developmental phenotypes are predictable from the earliest stages before visible aggregate formation begins. This predictability held across both genetic and environmental sources of variation, suggesting fundamental constraints on developmental trajectories and indicating that subtle phenotypic variations carry important information. These results demonstrate how machine learning can reveal hidden structure in complex multicellular dynamics and provide scalable methods for phenotypic analysis without manual annotation.

## Introduction

Multicellular self-organization is fundamental to biological processes across all kingdoms of life [5]. The development of a complex living system involves numerous iterative self-organizing steps, ranging from the formation of tissues and organs to the emergence of whole organisms and their interactions within communities. In bacteria, self-organization into structured biofilms can contribute to virulence and protect from environmental stressors such as desiccation or antimicrobial agents [39]. For example, developmental self-organization is an essential part of the bacterial stress response, as observed in the formation of starvation-induced fruiting bodies in the soil-dwelling bacterium *Myxococcus xanthus* [29].

*M. xanthus* serves as a model organism for studying the genetic basis of multicellular self-organization and its dynamics. Mutations in the *M. xanthus* genome can influence both individual cell behavior and multicellular dynamics, resulting in distinct multicellular phenotypes [29]. A central assumption in genetics is that mutations in genes within the same functional pathway will produce phenotypes with shared features, potentially distinguishable from those arising from mutations in unrelated pathways. If such phenotypic coherence is robust, persisting despite the confounding effects of pleiotropy and additional mutations, the motility systems of *M. xanthus* offer a natural test case. The bacterium employs two genetically distinct systems for gliding: adventurous (A) and social (S) motility [19]. Although their regulation intersects, the mechanics of A- and S-motility operate largely independently, such that mutations in the A-motility genes eliminate A-motility while leaving S-motility intact, and vice versa [46, 54]. Mutants within each system tend to exhibit distinguishable phenotypic features, raising the possibility that A- and S-system mutations leave persistent, if subtle, phenotypic signatures. However, in contrast to vegetative swarming assays, where defects in one of the motility systems can be diagnosed by examining the colony edge morphology [40, 34], no such approaches have been established for development.

Detecting these phenotypic signatures, especially when they are subtle, variable, or emerge gradually, requires methods capable of parsing the entire spatiotemporal dynamics of *M. xanthus* development. Time-lapse microscopy has become a powerful tool for recording these dynamics on agar substrates. Prior studies have used time-lapse imaging to describe and quantify swarm phenotypes at discrete timepoints, mainly in wild-type or mutant strains that reliably form aggregates [51, 2]. These approaches have provided valuable insights into the structure and timing of aggregate formation, but largely overlook early-stage swarm dynamics such as streaming, which may precede visible aggregate formation. Some mutant strains do not form aggregates, or produce only small, opaque, or transient ones that aggregate-focused methods have not attempted to characterize or compare across strains [31, 50, 28]. As a result, these methods do not capture important phenotypic features, both within immature aggregates and in the inter-aggregate space [23, 44]. Finally, since phenotypic features often emerge at different times across replicates or strains, comparisons based on fixed time windows may be misleading. Instead, robust analysis requires comparing full developmental trajectories.

Meeting this challenge requires computational methods that can quantify phenotypic similarity across entire time series without relying on aggregate parameters or annotated labels. Machine learning is particularly well-suited to this task because it can extract high-dimensional, temporally resolved features from complex image data without requiring predefined labels, explicit models of development, or spatial alignment. Traditional clustering methods, such as the nearest neighbor algorithm [9], rely on human-annotated labels for evaluating performance, often using metrics like precision (how accurately the model identifies true phenotypic matches) and recall (how completely it recovers all relevant phenotypes). These evaluation frameworks break down in label-free settings, where ground-truth annotations are unavailable.

Deep learning approaches have recently achieved remarkable success in biological image analysis, enabling more quantitative and scalable studies of cell and tissue dynamics [47, 6]. Convolutional and transformer-based models can automatically classify subtle cellular phenotypes from microscopy images and detect anomalies in medical scans with high accuracy [42, 35, 1]. At the same time, advanced segmentation networks can delineate individual cells or structures even in dense tissues and microbial communities, facilitating precise phenotypic quantification in complex biological systems [49, 16]. Beyond discriminative tasks, generative deep learning has opened new avenues in synthetic imaging [24, 20]. Diffusion models, for example, can enhance or predict high-resolution cellular morphologies from limited data [33]. Meanwhile, GAN-based frameworks, such as StyleGAN variants, have been applied to produce realistic microscopy images and even simulate time-lapse sequences that capture emergent collective behaviors [36]. Together, such state-of-the-art methods – spanning vision transformers, diffusion models, and GAN architectures – are reshaping biological imaging by enabling scalable, accurate, and versatile analyses that often surpass traditional feature-based techniques.

Despite these advances, current deep learning models face significant limitations in analyzing dynamic microbial behaviors, such as *M. xanthus* development. First, many architectures, particularly Vision Transformers, are architecturally dependent on absolute positional information, introducing spatial biases when the biological phenotype is position-invariant [4]. Second, models that focus on static end-point morphologies overlook the dynamical and transient features that define emergent self-organization, such as early streaming dynamics [23, 2]. Finally, a significant challenge lies in evaluation in a truly label-free setting [27, 48]. This reveals a critical disconnect: impressive performance on public benchmark datasets does not ensure that learned latent representations capture biologically meaningful features [12]. For our study, the primary objective is not to pursue a state-of-the-art generative model, but to obtain a robust latent space that faithfully represents cellular phenotypes—one that is low-dimensional, interpretable, and invariant to irrelevant positional variations.

To address these challenges, we developed an unsupervised deep learning framework that disentangles phenotype from position, enabling scalable, trajectory-level comparisons of *M. xanthus* development. To put this framework into practice, we first constructed a comprehensive 24-hour time-lapse microscopy dataset consisting of 937 movies from 292 genetically diverse *M. xanthus* strains (including wild-type and mutants), designed to capture a broad spectrum of phenotypes. Recognizing that existing models were either insufficiently robust, lacked interpretability, or were incomplete for this task, we designed our own framework. It integrates three key components: (1) a deep image encoder [17] to extract low-dimensional and interpretable latent features, (2) a generative image model [36] to ensure visually faithful phenotype reconstruction, and (3) a contrastive discriminator network [3] to enhance the robustness and consistency of the latent representations. This approach allows us to uncover hidden determinants of collective behavior from full developmental trajectories.

## Results

### Comprehensive Microscopic Image Dataset Reveals Temporal and Genotypic Variation in *Myxococcus xanthus* under Nutritional Stress

To generate a time-lapse microscopy dataset capturing the full scope of swarm phenotypes, we imaged both wild-type and mutant strains of *M. xanthus* over a continuous 24-hour period beginning shortly after the onset of starvation. The dataset was designed to capture a wide range of developmental dynamics by maximizing phenotypic diversity while minimizing variation introduced by experimental inconsistency.

We standardized sample preparation, microscopy conditions, and data acquisition protocols. We collected a total of 937 brightfield time-lapse movies representing 292 genetically distinct strains drawn from a collection colloquially known as the “Common Stocks” [29] (Methods Strains and Culture Conditions), with multiple replicates per strain (Table S1). Each movie consists of 1440 frames acquired at one-minute intervals, initiated within 15 minutes of starvation onset. The strains were selected to represent a broad phenotypic spectrum, allowing for a comparative analysis of diverse developmental trajectories.

Upon nutritional depletion, wild-type *M. xanthus* strain DK1622 gradually transitions from a dispersed distribution on agar surfaces to structured multicellular fruiting bodies [8]. In representative time-lapse images, distinct aggregates first become visibly apparent by 12 hours. From 12 to 24 hours, these aggregates progressively mature, exhibiting sharper boundaries, enhanced texture, and increased surface reflectivity (Figure 1A).

**Figure 1:**
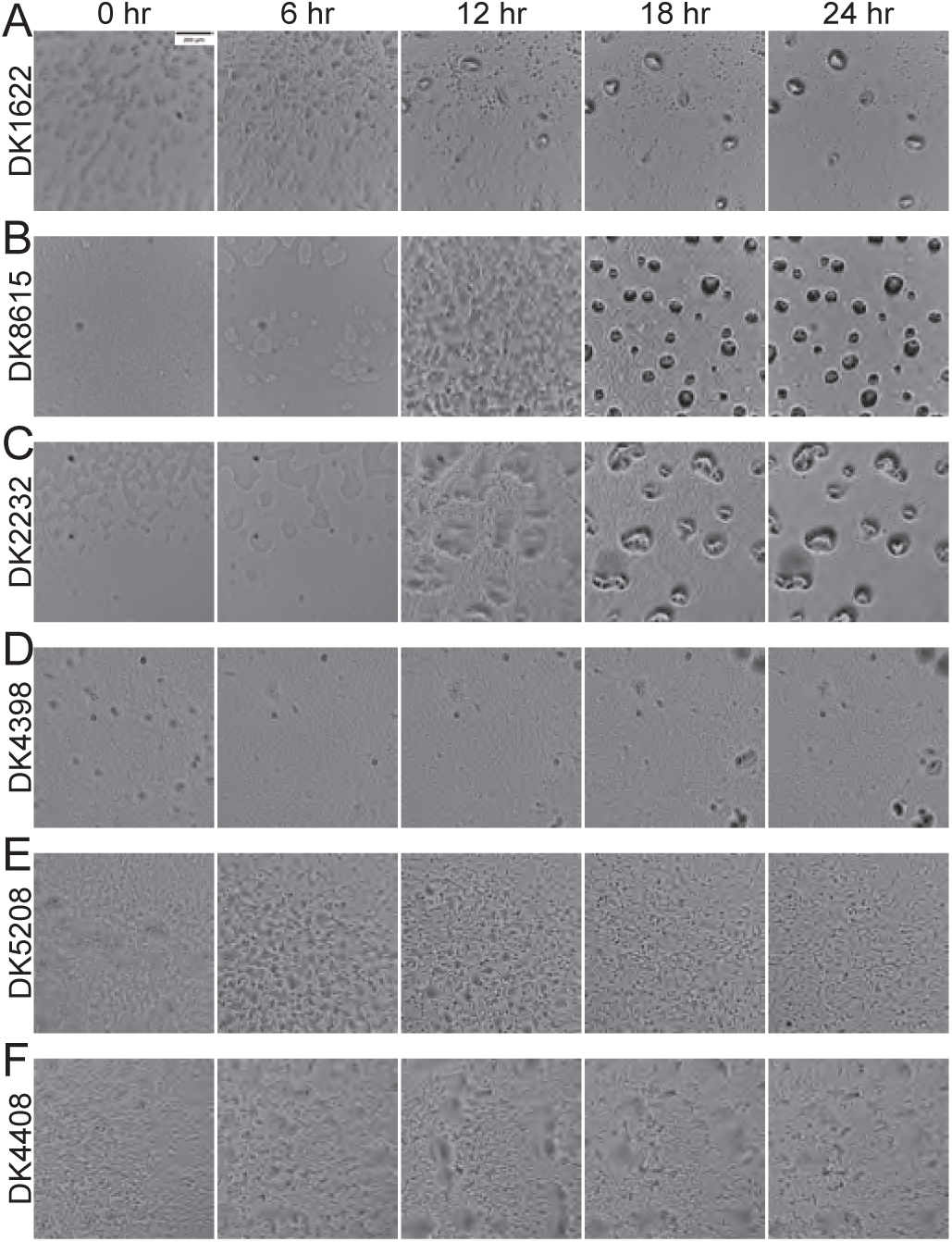
Differential Gene Effects on *Myxococcus xanthus* Emergent Behavior. (A-F) Cropped images of different strains captured at 0, 6, 12, 18, and 24 hours following nutritional stress. (A) Wild-type DK1622; (B) *pilQ* mutant DK8615, social motility impaired [11]; (C) *aglB3* mutant DK2232, adventurous motility impaired [14]; (D) *asgB* mutant DK4398, A-signal deficient [32]; (E) *csgA* mutant DK5208, C-signal deficient [30]; (F) *sdeK* mutant DK4408, sporulation deficient [15].

Genetic perturbations significantly alter developmental phenotypes by disrupting specific motility pathways. For instance, strain DK8615 [11], harboring a mutation in the *pilQ* gene, exhibits impaired S-motility, preventing aggregate formation at the 12-hour mark and yielding numerous thinner aggregates at 18 hours (Figure 1B). Conversely, strain DK2232 [14], carrying a mutation in *aglB3* crucial for A-motility, forms large, irregularly shaped, transparent aggregates starting at 12 hours, with increasingly pronounced morphological distortions by 18 and 24 hours (Figure 1C). Strain DK4398 [32], an *asgB* mutant lacking the A-signal necessary for intercellular communication, fails to initiate aggregate formation altogether (Figure 1D). Strain DK5208 [15], a *csgA* mutant deficient in the C-signal, initiates development—as indicated by subtle textural changes at 6 and 12 hours—but ultimately does not complete aggregate formation, even at later timepoints such as 18 and 24 hours (Figure 1E) [55].

The dataset also encompasses both extensively characterized and poorly understood genetic variants, offering opportunities to uncover new insights into phenotypes. For example, the *sdeK*-mutant strain DK4408, associated with a histidine kinase operating independently of A-signal, initially forms large, transparent, and irregular aggregates at 12 hours, resembling the *aglB3* mutant. However, unlike the *aglB3* mutant, these aggregates fail to mature further, maintaining their immature phenotype through 18 and 24 hours (Figure 1F) [15].

Phenotypic variation in *M. xanthus* arises from genetic mutations expressed across different stages of development following the onset of starvation, making precise and consistent phenotypic annotation challenging. Features such as immature aggregates with indistinct boundaries, transparent aggregates, variation in aggregate texture and reflectivity, and the presence of interconnecting streams between aggregates can be subtle, transient, or difficult to define precisely. Our dataset, acquired under standardized culture and imaging conditions, provides a valuable resource for studying this dynamic and heterogeneous phenotypic landscape.

### An Encoder-Generator-Discriminator Model Enables Deterministic Phenotypic Encoding

Quantifying the features and similarities of *M. xanthus* behavior without human-guided annotations requires machine learning directly from raw image data. To address this, we developed a model that combines three components: an encoder, a generator, and a similarity network. The encoder transforms each image into a numerical representation by producing a mean vector *µ* and a log-variance vector log *σ*^2^; from these, the feature vector *z* is sampled during training, while the mean vector *µ* is used during testing as a deterministic representation. A feature vector, *z*, together with a noise vector, *n* (sampled from a normal distribution), is used by the generator to reconstruct images that preserve phenotypic features of the input. To evaluate similarity, a Siamese network is used to compare real image pairs or real/reconstructed image pairs (Figure 2A, Table S2, S3, S4). This setup enables the system to capture phenotypic variation in an unbiased, category-free manner, without relying on human assumptions regarding what features are biologically meaningful. (Methods Network Architectures and Learning Algorithm)

**Figure 2:**
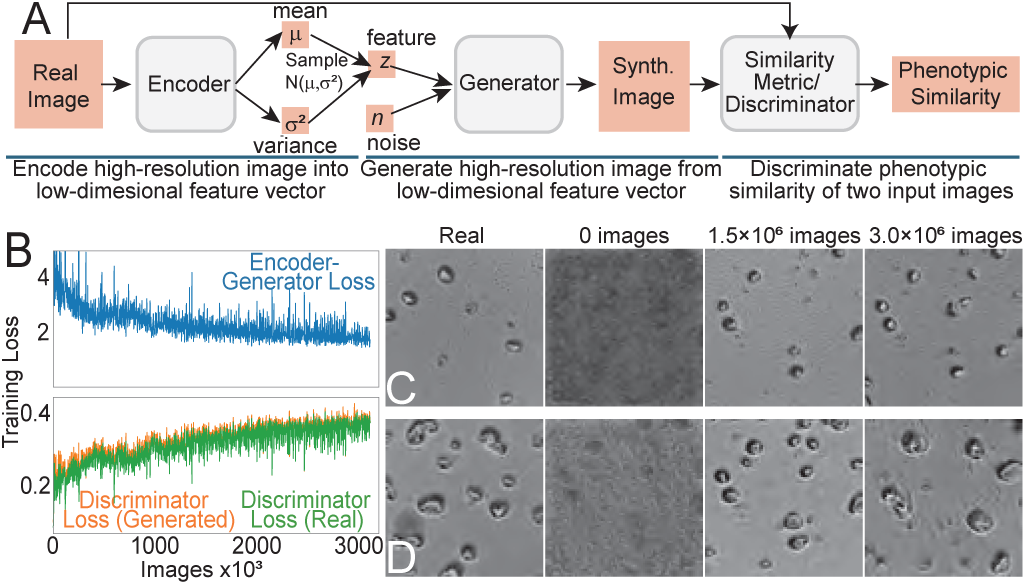
High-Fidelity Reconstruction of Microscopic Images Using a VAE-StyleGAN2 Integration. (A) Overview of the model architecture (details in Figure S1). (B) Training loss progression. Target image and model-reconstructed images for (C) a wild-type strain DK1622 and (D) A-S+ strain DK2232 at various training checkpoints.

The model was trained on 4 A40 GPUs using an unsupervised learning approach, processing approximately 3.0 × 10^6^ images over six days. Since the training was unsupervised, no validation set was used.

Training progress was monitored via loss curves, which plotted loss values against the number of processed images (Figure 2B). While all loss curves exhibited fluctuations, likely due to random noise and variations between image batches, the overall trends were clear. The encoder and generator losses steadily decreased, indicating the model’s improving ability to extract meaningful patterns and reconstruct realistic images. Conversely, the similarity network (discriminator) loss, which compares crops from real images with their reconstructions, showed a consistent increase. This trend suggests that the reconstructed images became progressively more realistic, making it more difficult for the discriminator to distinguish them from the originals. After six days of training, all loss curves had plateaued, indicating convergence and prompting the end of the training process.

The quality of the reconstructions, assessed by visual similarity to the input frames, improved throughout the training. Aggregates became more clearly defined, with increasingly accurate boundaries and internal textures. Initially, reconstructions appeared flat and lacked detail. After three days (approximately 1.5 × 10^6^ images), distinct aggregates began to emerge, though some texture details were still missing. By day six, the reconstructed aggregates closely resembled the target phenotypes (Figures 2C,D).

### Human Evaluation Confirms High Fidelity and Biological Alignment

To quantitatively assess the quality of model-generated images and their fidelity in reconstructing biologically meaningful phenotypes, we conducted two human evaluation studies using separate criteria for fidelity and alignment. Fidelity refers to how closely a model-generated image resembles a real microscopic image of *M. xanthus*, while alignment refers to how well a reconstructed image preserves the phenotypic characteristics of a specific input image. In both studies, participants were asked to indicate whether they were familiar with microscopic images and whether they had performed any experiments related to *M. xanthus*. In the analysis, participants who answered “yes” to both questions were grouped as experts, and the rest were grouped as non-experts. (Figure S3)

For the fidelity evaluation (Figure 3A), participants were asked to rate the fidelity of individual images using a 5-point Likert scale (1 = least realistic, 5 = most realistic). The image set consisted of 52 real microscopic images, 52 images generated by our model, and 52 images generated by StyleGAN2 [21]. These images were uniformly and randomly distributed across 13 pages. To monitor participant attentiveness, we included 13 synthetic images from StyleGAN3[22], which exhibit characteristic artifacts. As these images were expected to be rated poorly due to their visible artifacts, participants were considered attentive if they assigned a score of ‘1’ to at least one of them. Ratings from 18 experts and 21 non-experts met this criterion. Ratings were computed for each image and subsequently aggregated within each group. A one-way ANOVA followed by Tukey’s HSD test [45] and Hedges’ g effect sizes [18] were used to compare fidelity scores across image types. No statistically significant differences were found among real, our model, and StyleGAN2 images (mean ± SD: 3.3 ± 1.3, 3.5 ± 1.2, and 3.3 ± 1.3, respectively; all *p >* 0.5; all *g <* 0.1), indicating comparable visual realism. In contrast, StyleGAN3 images received significantly lower ratings (1.1 ± 0.5), with all comparisons yielding *p <* 0.001 and large effect sizes (*g >* 1.8), reflecting easily perceived artifacts (Figure 3C; Table S5). In this context, p-values assess statistical significance (*p <* 0.05 threshold), while Hedges’ g quantifies the magnitude of the difference, with *g >* 0.8 interpreted as large.

**Figure 3:**
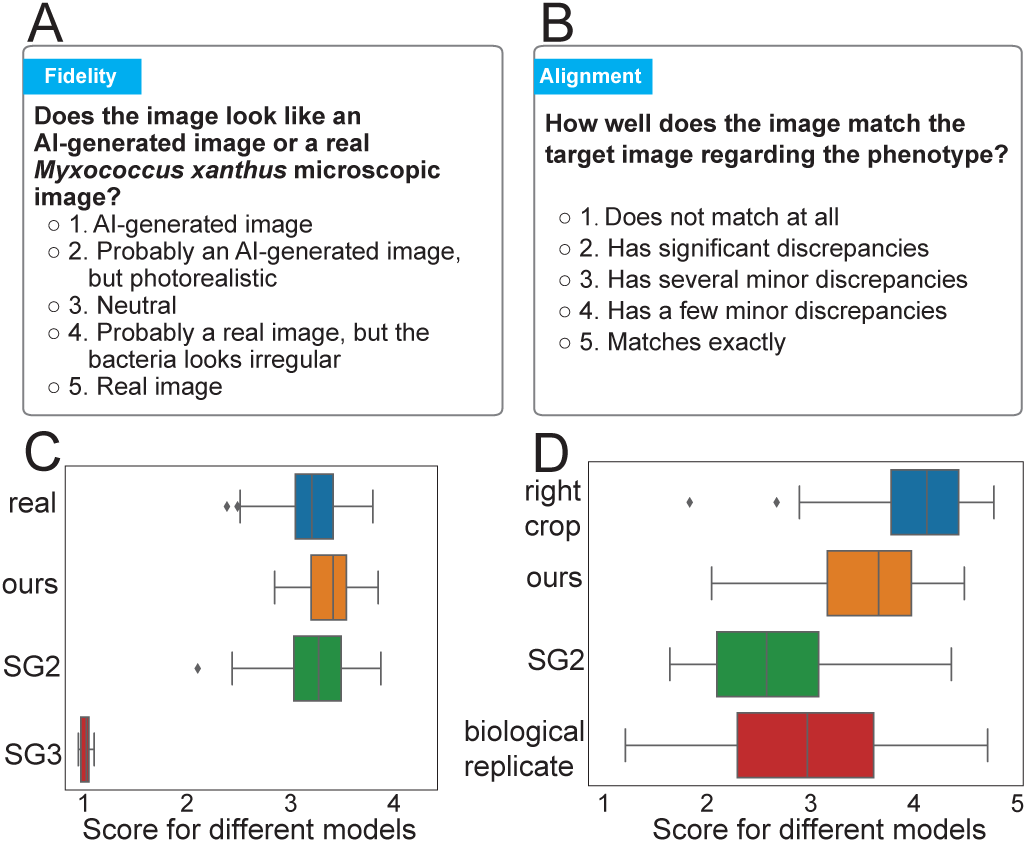
Human evaluation. (A) Question and labels of fidelity evaluation; (B) Question and labels of alignment evaluation; (C) Distributions of human ratings for fidelity; (D) Distributions of human ratings for alignment.

In the alignment evaluation (Figure 3B), each page presented a reference image, defined as the left square crop from a full microscopic view. Participants were asked to assign a similarity score to each of four target images (1 = does not patch at all, 5 = matches exactly), which included (i) the right square crop from the same microscopic view (“right”), (ii) the left crop from an experimental replicate of the same strain (“repeat”), (iii) an image reconstructed via latent-space back-projection using StyleGAN2 (“Style-GAN2”), and (iv) an image reconstructed by our model (“our”). A total of 50 reference images were evaluated. Tukey’s HSD test indicated statistically significant differences among all candidate groups. The right crop received the highest mean rating (4.1 ± 1.0), followed by our model’s reconstruction (3.6 ± 1.2). In contrast, the replicate and StyleGAN2 reconstructions received lower scores (2.9 ± 1.3 and 2.7 ± 1.2, respectively). Pairwise post hoc comparisons (Table S6) revealed that our model significantly outperformed both StyleGAN2 (p = 0.000, g = 0.70) and the experimental replicate (p = 0.000, g = 0.50), with medium-to-large effect sizes. Furthermore, the difference between our model and the right crop was statistically significant (p = 0.000, g = –0.45), indicating that adjacent regions from the same microscopic field were rated as more phenotypically similar than our reconstructions. Notably, Style-GAN2 reconstructions were rated significantly lower than experimental replicates (p = 0.000, g = −0.17), underscoring our model’s superior phenotypic consistency (Figure 3D; Table S6). Thus, our model can reconstruct realistically looking images that are closer to the reference image than the biological replicate of the same strain.

### Disentangled Phenotype and Position Enables Visualization of Phenotypic Trajectories

To investigate how our Variational Autoencoder represents image information, we reconstructed images by varying either the encoded feature vector *z* or the randomly sampled noise vector *n*, while holding the other fixed. This approach allowed us to examine the specific role of each vector in the reconstruction process. When we varied the input images, thereby changing the encoded feature vectors while keeping the noise vector constant, the reconstructions displayed clear differences in aggregate shape and organization. In contrast, positions of many aggregates appeared essentially unchanged (Figures 4B,C), suggesting that the feature vector captures phenotypic information (Figures 4A and 4B). On the other hand, when the feature vector was fixed and the noise vector was replaced with a different sample, the resulting image displayed a similar phenotype but with aggregates appearing at different positions (Figure 4C). These observations indicate that the model successfully separates phenotype-related variation (captured in the feature space) from positional variation (captured in the noise space).

**Figure 4:**
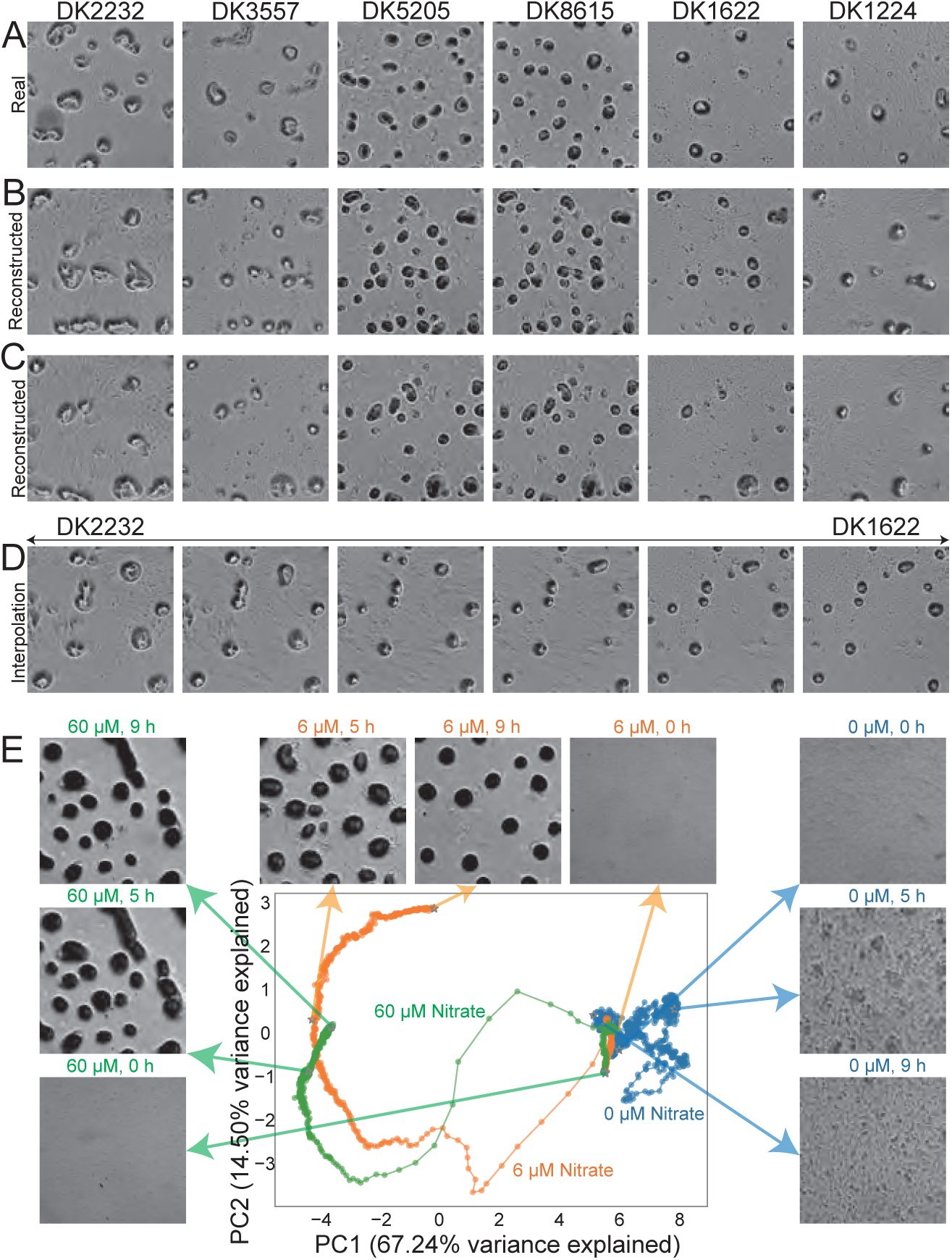
Image Reconstruction. (A) Target images; (B) Reconstructed images with fixed noise; (C) Reconstructed images with another fixed noise; (D) Linear interpolation in the feature space between the encoded image of DK2232 and DK1622; (E) The PCA-processed encoded movie trajectories of LS3934 from Shimkets’ dataset. Selected frames are shown aside.

To further evaluate whether the feature space changes smoothly across different phenotypes, we selected two target images exhibiting distinct phenotypes and encoded them into feature vectors corresponding to strains DK2232 and DK1622. We then reconstructed images by linearly blending between these vectors while keeping the noise input fixed (Figure 4D). The interpolated reconstructions reveal a gradual transformation in aggregate morphology, with aggregates shrinking and separating progressively. This smooth transition highlights the continuity of the feature space, providing an intuitive visualization of the phenotypic differences between the two strains.

We also tested the model’s ability to generalize to new experimental conditions and to track phenotypic changes over time and over conditions. For this purpose, we utilized an independent dataset from Larry Shimkets’ lab, which was acquired under different growth conditions, lighting, and resolution (Methods Time-Lapse Imaging Shimkets Lab). After normalizing the images for brightness and resolution, we focused on a subset from strain LS3934, which expresses a modified version of the DifA chemoreceptor (NafA) [53], where the sensor domain is replaced by a nitrate sensor. We applied principal component analysis (PCA), a method that reduces high-dimensional image features into a few orthogonal axes capturing the dominant sources of variation, and defined a principal component (PC) space using 21 time-lapse movies sampled at 60-minute intervals. We then encoded the 0-9 hour images from movies obtained under varying nitrate concentrations (0, 6, and 60 *µM*) by projecting them into this PC space.

Figure 4E (Supplementary Movie 3) shows the trajectories of these movies projected into a two-dimensional space defined by PCA. In the absence of nitrate (0 *µM*), aggregates do not form, and the trajectory remains relatively static. At a concentration of 6 *µM* nitrate, aggregates begin to form gradually, resulting in an extended trajectory along the first principal component (PC1). By 5 hours, distinct, high-density aggregates emerge, followed by a reduction in aggregate size and the disappearance of some aggregates between 5 and 9 hours. Under a higher nitrate concentration (60 *µM*), aggregate formation follows a similar temporal pattern; however, the aggregates are significantly larger. This is reflected by a lower value along PC2 at 5 hours and a final position at 9 hours that resembles the 5-hour state under 6 *µ*M. These results demonstrate that our model not only generalizes to out-of-distribution datasets but also captures the evolving phenotypic landscape over time.

### Model Features Preserve Intra-Image Consistency in an Unsupervised Setting

To evaluate the relevance and consistency of the learned features without using labels, we performed a simple test using two subregions from each image. Specifically, we selected two fixed, non-overlapping crops (each 512 × 512 pixels) from the left and right parts of each image. From each crop, we extracted features using three methods: (i) raw pixel values, (ii) 22 human-defined features, and (iii) 13 features from our unsupervised model. As a baseline, we also compared crops randomly paired from different images (Figure 5A).

**Figure 5:**
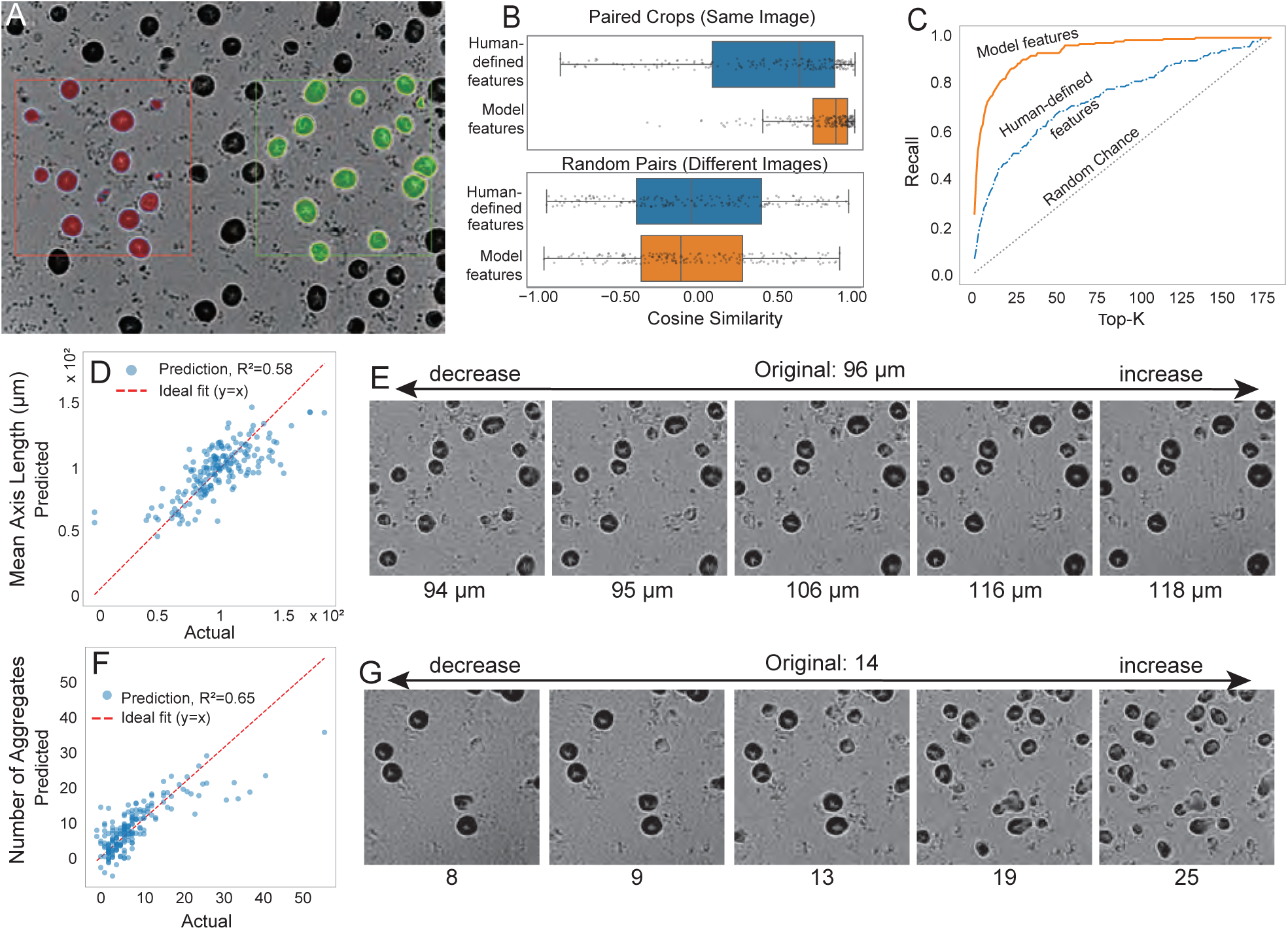
Evaluation and interpretation of phenotypic features. A) Image processing workflow. Two fixed, non-overlapping crops (left and right, each 512 × 512 pixels) were selected from each image (original size: 1296 × 972 pixels). Aggregates were segmented as described in the methods section and assigned to either the left or right crop based on the location of their centers. Edges of left and right crops, as well as the assigned aggregates, are colored in red and green, respectively. B) Cosine similarity analysis. Raw pixel intensities, human-defined features, and model-extracted features were normalized and reduced to their first five principal components (PCs) based on left crops. Box plots compare cosine similarities between paired left-right crops from the same image (top) versus randomly matched pairs (bottom). C) Retrieval analysis. Recall curves indicate the percentage of correctly retrieved matching right crops ranked within top-K nearest neighbors for each feature set based on PC1–5. D, F) Linear regression predicting total aggregate area (D) and number of aggregates (F) from model-extracted features, with performance metrics (D: *R*^2^ = 0.68, mutual information=0.12; F: *R*^2^ = 0.65, mutual information=0.09). E, G) Generated images showing phenotypic variation along the regression weight vector directions obtained from analyses in (D) and (F), respectively, starting from the feature vector of the left crop shown in panel (A).

We expected crops from the same image to be more similar to each other than crops from different images.

If the learned features effectively capture the visual characteristics of the image, they should exhibit a stronger similarity between the two crops. Because raw feature sets often contain strong correlations that can artificially inflate cosine similarity, we first applied z-score normalization to each feature type and then performed Principal Component Analysis (PCA) on the features of the left crop. Retaining only the top five principal components ensured that similarity was assessed using the dominant, non-redundant sources of variation rather than noisy or collinear dimensions. These principal components were then used to project both left and right crops, after which cosine similarity was calculated between each pair of crops. This metric quantifies how similar two vectors are by measuring the cosine of the angle between them, ranging from 1 (identical direction) to 0 (orthogonal, no similarity) to –1 (opposite direction) [38].

As shown in Figure 5B, all feature types showed higher similarity between two crops from the same image than between randomly matched crops. Model-extracted features showed the strongest consistency, outperforming human-defined features and raw pixels. This result suggests that the model captures meaningful spatial patterns even when no obvious structures, such as aggregates, are present, which human-defined features often overlook. To further illustrate this point, additional examples in the supplementary figures illustrate cases where the model features correctly match crops while the human-defined features do not (Figure S4).

To assess how well different feature representations capture phenotypic similarity, we designed a crop-matching task. For each image, we assumed that the left and right crops depict the same phenotype. Using the left crop as a query, we ranked all 184 right crops by cosine similarity in feature space. We then asked whether the true matching right crop was found within the top-K most similar candidate images. Plotting this success rate across different values of K yields a recall–top-K curve, with the area under this recall-top-K curve (AUC) summarizing the overall retrieval performance across different K values. The AUC ranges from 0.0 to 1.0, where 1.0 represents a perfect ranking (i.e., the true match is consistently ranked at K=1), and a score approaching 0.5 indicates random-chance performance.

The top five PCs of model-extracted features achieved the highest performance (AUC ≈ 0.94), substantially outperforming human-defined features (AUC ≈ 0.75) and raw pixel intensities (AUC ≈ 0.70) (Figure 5C). These results demonstrate that model-derived features provide a more reliable measure of phenotypic similarity than traditional approaches.

To interpret model-derived features in terms of human-defined metrics, linear regression was performed to predict selected human-defined features (total aggregate area and aggregate count) from the 13-dimensional latent features. These metrics were selected due to their relatively high predictive performance, yielding coefficients of determination (*R*^2^) of 0.68 and 0.65, respectively (Figure 5D, F). Nonetheless, these correlations indicate only moderate linear relationships.

To visually interpret the latent feature space, we extrapolated phenotypes along the directions of the regression weights (Methods Feature Correlation Analysis). Starting from the central reconstructed image (0 std), we varied latent features by ±2 and ±4 standard deviations (std). Extrapolating along the regression direction for total aggregate area resulted in increased aggregate sizes and counts for positive extrapolation and decreased values for negative extrapolation (Figure 5E). Conversely, extrapolating along the aggregate-count direction increased aggregate numbers but reduced individual aggregate sizes (Figure 5G). Negative extrapolations (−4 and −2 std) along the aggregate-count direction showed minimal phenotypic change, suggesting a nonlinear relationship between latent features and human-labeled metrics.

Crucially, the robustness of these learned features extends the scope of quantitative analysis to scenarios where human-defined features fail. For instance, our model successfully extracts features from early-stage movie frames before aggregates are well-formed, or from genetic mutations that prevent the formation of mature aggregates. By doing so, our model not only increases accuracy but also expands the coverage of quantitative phenotypic description.

### The model identifies developmental signatures of motility mutants

We next ask whether the feature space encoding of microscopic images can help group mutants corresponding to the same pathway by capturing subtle phenotypic information. To address this question, we first sought to classify strains with distinct genetic defects in motility, specifically A motility-deficient (A-S+) versus S motility-deficient (A+S-) [41], using their 13-dimensional feature trajectories. To this end, we utilize the strain genotype annotation to identify 45 movies from 14 strains labeled A+S- and 44 movies from 15 strains labeled A-S+ in our time-lapse imaging dataset (Table S1).

Each image at a given time point was represented by a 13-dimensional feature vector. We first focused on the final time point (24 h), where phenotypes were highly variable; some strains from both genotypes formed aggregates, while others failed to do so (Figure S5B). Despite this phenotypic heterogeneity, we tested whether the underlying genotype could be predicted from the feature vectors. We trained a Support Vector Machine (SVM) model [7] using a 3-fold cross-validation strategy [25] grouped by strain to ensure generalization. Model performance was evaluated using the receiver operating characteristic (ROC) curve and its area under the curve (AUC) [13], which measures the model’s ability to correctly rank samples from different classes, regardless of the decision threshold. A ROC-AUC of 1 indicates perfect discrimination, whereas 0.5 corresponds to random guessing. We report the mean and 95% confidence interval (CI) of the ROC-AUC, where the CI represents the range of values expected to contain the true performance 95% of the time if the experiment were repeated. Under this framework, the model achieved a ROC-AUC of 0.85±0.04 (mean ±95% CI) at 24 h, demonstrating robust genotype discrimination even in the presence of phenotypic variability. More surprisingly, repeating this analysis for the initial time point (0 h) revealed that the genotypes were separable even from the earliest moments of observation, with a ROC-AUC of 0.91±0.06 (Figure 6A). Visualization of the PCA space confirmed a clear separation between the two classes at both time points, demarcated by the SVM-derived decision boundary (Figure 6B). This result indicates that phenotypic differences associated with A- versus S-motility defects are encoded in the latent feature space from the very onset of development, well before aggregates form and motility differences become visually apparent.

**Figure 6:**
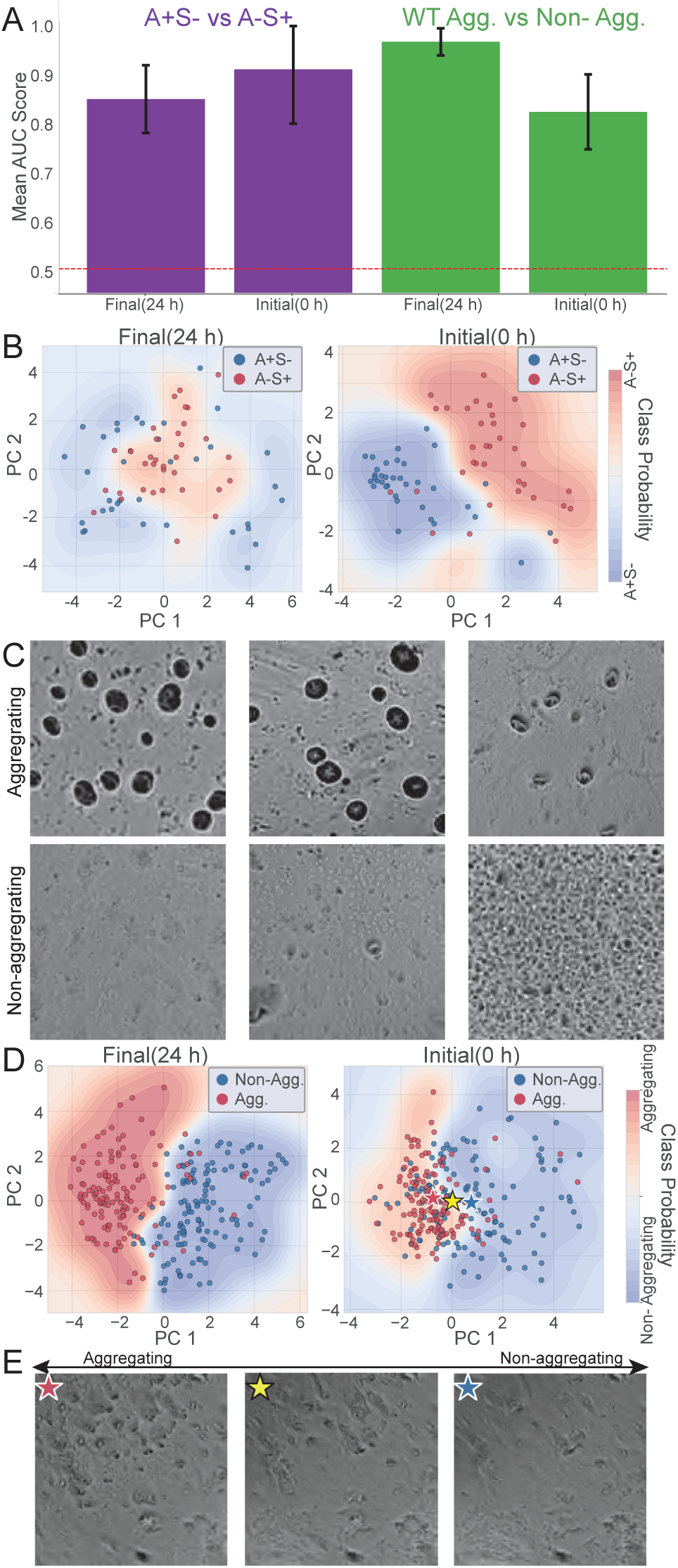
Latent space features allow for predicting motility genotype and aggregation fate. (A) ROC-AUC scores show high classification accuracy between A+S- and A-S+ and between aggregating and non-aggregating WT conditions at both final and initial time points. (B) At both 24 h and 0 h, the populations are separable in the PCA feature space for motility mutants. Contours are SVM-derived class probabilities. (C) Different 24 h images of WT. Top: aggregating; bottom: non-aggregating. (D) At both 24 h and 0 h, the populations are separable in the PCA feature space for WT aggregation. Contours are SVM-derived class probabilities; stars are class centroids. (E) Reconstructions from 0 h centroids visualize the subtle phenotypic features—imperceptible to the human eye—that drive the model’s accurate prediction.

### Latent Space Features Reveal Early Determinants of Developmental Fate

Having established that the model can distinguish genetically distinct strains, we next investigated whether it could detect more subtle phenotypic variations within a genetically uniform sample set. To examine the range of phenotypic outcomes in wild-type *M. xanthus*, we analyzed 352 movies prepared under the standard developmental protocol. We observed substantial variation in developmental outcomes, with a significant fraction of the movies failing to form visible aggregates by 24 h (see Figure 6C for representative examples). This unexpected heterogeneity provided an opportunity to test whether early developmental features could predict ultimate developmental fate.

We manually classified each movie as “Aggregating” (220 movies) or “Non-aggregating” (132 movies) based on whether visible aggregates formed by 24 h (Figure 6C). Using these labels, we trained an SVM classifier to test whether developmental fate could be predicted from the extracted feature vectors. We employed a 3-fold cross-validation grouped by experimental replicates to ensure robust generalization across independent experiments.

As expected, an SVM classifier easily distinguished between these outcomes using the 24 h feature vectors, achieving a ROC-AUC of 0.97±0.01. The score was not a perfect 1.0, likely reflecting the continuous nature of development, where some strains were in intermediate aggregation states in manual annotation. More strikingly, the classifier also achieved high performance on the initial T=0 h frames, yielding a ROC-AUC of 0.82±0.04 (Figure 6A). The separation between the two fate groups is readily apparent in the PCA space at both time points (Figure 6D).

This high predictability from initial time points strongly suggests that the ultimate developmental fate is encoded in subtle patterns present at the onset of starvation. To visualize these cryptic features, we reconstructed images from the mean T=0 h feature vectors for the “Aggregating” and “Non-aggregating” classes. Linearly interpolating between these two centroids reveals that the model identifies faint patterns of higher local cell density and early cell alignment as key predictors of future aggregation (Figure 6E). While consistently captured by the model, these features are often obscured by noise and spatial heterogeneity in individual raw images, making them nearly imperceptible to a human observer (Figure S5A).

## Discussion

In summary, we have demonstrated that, following training, the developed model architecture can map microscopy images into a low-dimensional feature space and utilize these features to generate realistic images that capture the phenotypic variation of *M. xanthus* developmental patterns. The resulting lowdimensional representation can then be used to quantify subtle changes in self-organization patterns, compare developmental trajectories of different strains, and even predict the developmental outcomes.

By representing developmental movies as trajectories through latent space, our framework enables new forms of quantitative phenotypic analysis that extend beyond static classification. This approach allows us to compare temporal dynamics across strains, detect and characterize different types of outliers, and identify divergence points associated with developmental bifurcations. Furthermore, by training simple regression models on latent features, we can predict human-defined phenotypes such as aggregate area or count, and identify latent directions that correspond to interpretable axes of variation. In some cases, extrapolating along these directions allows us to visualize how specific phenotypes emerge or are suppressed, offering a generative perspective on developmental variation. These findings underscore the utility of latent trajectory representations for phenotypic inference, dimensionality reduction, and the generation of mechanistic hypotheses in developmental systems.

Beyond identifying signatures of specific genetic perturbations, the model successfully distinguished motility genotypes (A+S- vs A-S+) without supervision, demonstrating its capacity to discover genetic structure within phenotypically complex populations. The Common Stocks strain collection, assembled over three decades of classical mutagenesis studies, contains many multi-mutant strains with secondary mutations introduced into backgrounds already deficient in motility or signaling pathways. These strains are phenotypically and genotypically complex, with poorly characterized developmental profiles and partially undocumented mutational histories, making it nearly impossible to infer genetic relationships solely from visual inspection. Remarkably, our unsupervised model identified phenotypic coherence among strains sharing functional defects A-motility and S-motility systems, even without curated genetic annotations.

Although A- and S-motility are mechanistically distinct systems for surface translocation, with their phenotypic differences most apparent during vegetative motility [40]. Nevertheless, our results indicate that strains carrying mutations in these pathways are distinguishable with high classification accuracy at both 24h and 0h. This result demonstrates that phenotypic differences associated with motility system defects are encoded in the latent feature space from the earliest observable time point. The basis for this early separability likely reflects pleiotropic effects of mutations in motility genes on cellular properties beyond locomotion itself, such as cell surface adhesion molecules, pili expression patterns, or exopolysaccharide production, that influence how cells initially settle and organize on agar surfaces [26]. These findings suggest that motility mutants should be understood not simply as movement-deficient but as having broader alterations in collective organization that manifest even in the absence of active translocation. More generally, the early distinguishability of genetically distinct strains, even when their defining phenotypes emerge only later, underscores the model’s capacity to extract subtle signatures of genetic perturbations from developmental trajectories.

Even *M. xanthus* wild-type populations do not invariably succeed in forming aggregates or fruiting bodies [37]. When analyzing 352 wild-type movies prepared under standardized protocols [43], we observed that 132 failed to form visible aggregates by 24h. Notably, our model could reliably distinguish between successful and failed developmental trajectories from the earliest stages of starvation. These failures clustered by experiment, with movies prepared on the same day from the same agar batch and culture stock showing similar outcomes. Admittedly, culture parameters do vary within conventionally acceptable ranges—optical density at harvest (Klett, 80-120), liquid culture incubation time (16-24 hours), and agar plate age (3-5 days)—as these kinds of laboratory day-to-day variations are typically considered physiologically irrelevant. Whether these or other unmeasured variables (such as agar batch composition, ambient temperature, and humidity) account for the observed clustering remains uncertain. However, regardless of the underlying cause, the clustering of developmental outcomes by experiment and the model’s ability to predict these outcomes from T=0 suggest that developmental fate may be more contingent on pre-starvation conditions than previously recognized.

The ability to predict developmental outcomes from the earliest timepoints indicates that fate-associated information is present well before aggregates become visible. Visual inspection reveals no apparent differences between aggregating and non-aggregating populations at T=0 hours (Supplementary Figure 5A), yet the model consistently extracts distinguishing features. The interpolation between class centroids (Figure 6E) suggests these features include subtle variations in local cell density and spatial organization. Whether these early spatial patterns directly influence developmental progression through biophysical mechanisms or serve as biomarkers of pre-existing cellular or environmental differences that independently affect both initial settling and subsequent development remains to be determined.

Regardless of the mechanisms, the factors operating at or before the onset of starvation play important and previously unrecognized roles in shaping developmental outcomes. Traditionally, the time prior to the formation of streams and nascent aggregates was treated as a relatively featureless “preparatory phase”, with developmental fate shaped afterwards, primarily by signaling cascades and transcriptional programs that unfold over subsequent hours [10]. Our results challenge this view, demonstrating that distinguishing information is present within minutes of starvation and that factors operating at or before this critical transition constrain subsequent trajectories more strongly than previously recognized. These results suggest that understanding developmental robustness will require a closer examination of how populations transition from vegetative growth to developmental commitment.

More broadly, this work demonstrates that machine learning approaches can reveal temporal structure in developmental processes that remains invisible to human observation, providing a complementary strategy to hypothesis-driven genetics for discovering when and how fate becomes determined in complex multicellular systems. Even though this study focuses on *M. xanthus*, the approach is broadly applicable to other biological systems in which dynamic two-dimensional patterns unfold over time and can be captured by time-lapse microscopy. The model does not rely on organism-specific features or supervised labels, making it adaptable to other biofilms, multicellular tissues, or synthetic systems in which morphology and dynamics encode functional information. The approaches developed can extract latent phenotypic structure, temporal dynamics, and developmental logic from complex image datasets. As imaging datasets continue to grow in size and diversity, such models may serve not only as tools for phenotypic classification but also as generative frameworks for mechanistic discovery and systems-level understanding.

## Materials and Methods

### Strains and Culture Conditions

To minimize phenotypic bias in strain selection, mutant strains were mainly chosen at random from the Common Stocks collection, a duplicate of the original collection housed at UC Davis (M. Singer, personal communication) and maintained at −80°C at Syracuse University. Wild-type strains were intentionally excluded to focus the dataset on mutant phenotypes. Selected strains were streaked onto CYE agar plates to verify culture purity before use. For time-lapse imaging, strains were grown at 32°C in CYE broth with shaking to mid-log phase, then processed as described below. Data collection prioritized generating a large, diverse dataset for model training rather than achieving uniform replication across all strains. Movies were acquired continuously, and experiments that failed due to technical issues (e.g., computer crashes, agar desiccation) were replaced with new strains rather than repeated to maintain consistent replicates. This approach yielded 937 movies from 292 genetically distinct strains, with variable replication (up to 3 replicates per run, and up to 5 runs per strain). Strains retained their original Common Stocks designations, and phenotypic annotations were obtained from the Common Stocks database (Table S1) and primary literature.

### Time-Lapse Imaging Welch Lab

Time-lapse microscopy was performed using a cluster of up to 96 custom-built 3D-printed microscopes [23], each equipped with a 4× objective lens providing 40× total magnification, a Raspberry Pi camera, red-light brightfield illumination, and a temperature-controlled stage maintained at 32°C via a Peltier device. All microscopes were networked via Ethernet to a central hub computer running PiServer software, enabling simultaneous control and centralized data storage. Custom Python software controlled image acquisition via SSH.

To induce development, cells from long-term stock cultures were streaked onto CYE agar and grown in CYE broth with vigorous shaking at 32^◦^C to a density of 4 × 10^8^. Cells were pelleted by centrifugation, washed, and resuspended in TPM buffer [10 mM Tris (pH 7.6), 1 mM KH_2_PO_4_, 8 mM MgSO_4_] at a final concentration of 4× 10^9^ cells/mL. 10-*µ*L spots containing approximately 4 × 10^7^ cells were deposited onto TPM agar slides [10 mM Tris·HCl, 8 mM MgSO_4_, 1 mM K_2_HPO_4_-KH_2_PO_4_, 1.5% Difco agar, pH 7.6] and allowed to dry before imaging. Images were acquired at one-minute intervals for 24 hours, yielding 1440 frames per movie.

### Time-Lapse Imaging Shimkets Lab

Time-lapse microcinematography was conducted as previously described [52] with the following modifications. *M. xanthus* strain LS3934 produces the interspecies chimeric protein NarX-DifA, known as NafA) [53]. LS3934 cells were grown at 32°C in casitone-yeast extract (CYE) broth (1.0% Bacto Casitone, 0.5% Difco yeast extract, 10 mM 3-[N-morpholino] propanesulfonic acid, pH 7.6, 0.1% MgSO4) with vigorous shaking. To induce development, cells grown to a density of 5 × 10^8^ cells/mL in CYE broth were pelleted by centrifugation and resuspended at a density of 5 × 10^9^ cells/mL in sterile distilled water; 20-*µ*L spots were plated on tris-phosphate-magnesium (TPM) agar (10 mM Tris·HCl, 8 mM MgSO4, 1 mM K2HPO4-KH2PO4, 1.5% Difco agar, pH 7.6) or TPM agar supplemented with KNO3 and allowed to dry. NO_3_ binds to the NarX sensor domain of NafA instead of the endogenous DifA signal to activate the Dif signaling pathway, thereby stimulating LS3934 development in a concentration-dependent manner. Cell spots were filmed using a Wild Heerbrugg M7 S dissecting microscope at room temperature. Photographs were taken every 5 min as .jpg files with a Spot Insight 2 camera using SPOT software v4.5 (Diagnostic Instruments).

### Image Processing

The original images, stored as JPEG files, have a resolution of 2592 × 1944 pixels, with a pixel size of 1 µm/pixel, and a file size of approximately 4.3 MB each, resulting in a total memory usage of 8.54 TB. Meanwhile, there are significant grid artifacts on 4 edges of the images. To address the large memory usage and eliminate grid artifacts visible at image boundaries, we first applied a Gaussian blur using a 5×5 kernel to smooth out grid artifacts. Then we resized images from a resolution of 1 µm/pixel to 2 µm/pixel, reducing the dimensions to 1296 × 972 pixels. After preprocessing, the images were saved in JPEG format with a compression quality of 60, resulting in a significantly reduced file size of approximately 126 KB per image. This adjustment decreased the total memory usage to 224.12 GB.

To enhance the diversity of training data and simulate different imaging conditions, in the training set, two regions from the same image were randomly cropped and rotated to generate images with identical phenotypes but varying positional information. For the validation set, cropping parameters were recorded and consistently applied to ensure reproducibility of the image set across different validation runs. For the distance calculation, the brightness of images was normalized to a mean value of 107.2. This standardization is crucial as brightness variation can significantly affect the calculated distances in feature space, impacting model performance.

### Dataset Splitting

To ensure class balance during training, we employed stratified sampling using rough phenotype labels (details provided in Supplementary Result 1).

For the image reconstruction tasks, one movie from each of 100 distinct strains was initially designated as the human evaluation test set. However, due to constraints on questionnaire length, a subset comprising 50 strains was ultimately reported.

### Network Architectures and Learning Algorithm

To assess phenotypic similarity, we design a Siamese network that processes image pairs through identical ResNet backbones. Each branch outputs a 4 × 4 feature map; these are concatenated and fed into a CNN to yield a scalar similarity score. For two random crops from the same microscopic view, **x**_1_ and **x**_2_, and a generated image defined as

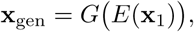

The discriminator loss is decomposed into four sub-losses:

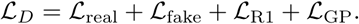

where *L*_real_ is the real pair loss, *L*_fake_ is the fake pair loss, *L*_R1_ is the R1 regularization, and *L*_GP_ is the gradient penalty. See supplementary for details.

For image synthesis, the VAE encoder (based on ResNet) maps an input image into the mean and log-variance of an *n*-dimensional latent code. The encoder outputs *µ* and log *σ*, from which a latent vector is sampled. The generator is composed of two parts: first, a mapping network transforms the sampled latent code into a higher-dimensional intermediate latent vector **w**; then, a synthesis network uses **w** to modulate the generator’s weights to produce the final image, following the inherent StyleGAN2 design. The overall loss for the encoder–generator branch is then summarized as:

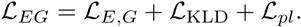

where *L_E,G_* is the adversarial loss, *L*_KLD_ is the KL-Divergence loss, and *L_pl_* is the path length penalty. See supplementary for details.

#### Training Schedule

The VAE and similarity metric are trained in an adversarial manner. In each training round, the discriminator is updated for three iterations, followed by one update of the encoder–generator branch. This alternating schedule ensures that the generated images gradually learn to “fool” the discriminator while maintaining a well-structured latent space. All networks are optimized using the Adam optimizer with a learning rate of 1 × 10^−3^.

### Parameter Initialization

To remove positional information from the feature representation, we initially trained our model on brightfield microscopic images of *Myxococcus Xanthus* from Larry Shimkets’ lab[51], adjusted to approximately 2 µm/pixel from 2.3 µm/pixel using a scale factor of 1.15. From this dataset, we extracted one image every 30 minutes, resulting in a total of 16,166 frames from 893 movies. The model, pretrained on 302k images, showed optimal separation of positional data at this stage. Further training was conducted on the Welch’s dataset, avoiding later stages where positional information re-enters the feature space. The model was trained on 4 NVIDIA A40 GPUs with a batch size of 112.

### Human Evaluation

#### Fidelity Assessment

We assessed the fidelity of the images using a 5-point Likert scale to determine if the images were AI-generated or real. The evaluation involved 30 real image crops, 30 images generated by our model, 30 by the StyleGAN2 model, and 10 by the StyleGAN3 model.

#### Alignment Assessment

To evaluate phenotypic alignment, a target image was compared against four options using a 5-point Likert scale. This test utilized the left crop of 30 real images, corresponding right crops, their experimental replicates, and images reconstructed by our model and the StyleGAN2 model.

### Trajectory Visualization

To visualize the evolution of the 13-dimensional features extracted from the selected movies, we applied principal component analysis (PCA) to reduce the feature space. The 13-dimensional trajectories were then projected onto the first two principal components, allowing us to capture and visualize the dominant patterns of temporal change.

### Feature Comparison

To evaluate the effectiveness of model-derived representations in capturing phenotypic features, we compared them with manually defined features extracted from segmented aggregates using the method described in [51]. Each image was cropped into left and right regions, and both manual and model-based features were computed.

To ensure segmentation quality, we applied empirically chosen thresholds to filter out images with abnormal or noisy segmentations. Images were excluded if fewer than half of the aggregates were larger than 100 pixels (resolution: 1.62 µm/pixel) or had eccentricity*<* 0.8, or if any aggregate exceeded 100,000 pixels or a major-to-minor axis ratio *>* 3. These thresholds were selected to remove extreme outliers based on visual inspection and prior experience, rather than to define biological phenotypes. A total of 184 images passed this quality control step.

For each image, we computed a set of human-defined features, including the mean, min, max, median, and standard deviation of area, perimeter, eccentricity, and axis length, along with the total area and object count, and our model-extracted features. Both feature sets were z-score normalized. We then performed a retrieval task: images were split into left and right crops. Principal Component Analysis (PCA) was applied to the left-crop features, and the top 5 principal components (PC1–5) were used to compute cosine similarity between left and right crop vectors. Retrieval curves were generated by ranking the similarity of each right crop to all left crops and recording the rank of the correct match, comparing paired (correct) vs. random pairings.

### Feature Correlation Analysis

To interpret the 13-dimensional feature vector *z* of our model, we trained a linear regression model to predict each human-defined feature *f* (*z*) from *z*, using the form:

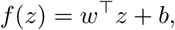

where *w* ∈ R^13^ is the weight vector and *b* is the bias term.

To visualize how changes in human-defined features affect the generated images, we perturbed the encoded feature *z* along the direction of the regression weights *w*, which correspond to the gradient 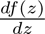. Specifically, we modified *z* according to:

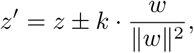

where *k* is a scaling factor controlling the magnitude of the change in *f* (*z*). This formulation ensures unit effect-size scaling, where 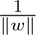 cancels the effect of coefficient magnitude, allowing consistent interpretation of the directionality.

By reconstructing images from the perturbed feature vectors *z*^′^, we generated synthetic phenotypes that systematically vary along interpretable biological dimensions. This analysis provides a direct, visual interpretation of how latent representations in our model relate to specific phenotypic traits.

### Classification and Performance Evaluation

We trained Support Vector Machine (SVM) classifiers on the 13-dimensional feature vectors at 0 h and 24 h to distinguish aggregating versus non-aggregating phenotypes. Model performance was evaluated using stratified cross-validation, with strain-level grouping to avoid data leakage when analyzing motility mutants. Classification accuracy was quantified by the ROC-AUC. For visualization, feature vectors were projected into PCA space and the SVM decision boundary was displayed in 2D. See supplementary for details.

## Acknowledgments

We thank Anke Treuner-Lange, David Zuckerman, Qiwei Li, Emilia MF Mauriello, Lee Kroos, Daniel Wall, Christopher Vassallo, Xin Luo, Xingwen Chen, Zhaoyang Zhang, Ting Lou, Erdong Ding, Tripti Midha, Liudeng Zhang, and 25 other anonymous participants for taking part in our human evaluation study survey. The authors are also grateful to Chris Cotter for the discussion and ideas that led to the initiation of this project. This research was primarily funded by NSF IOS awards 1856742 and 1856665, with partial support from NSF IOS-1951025.

## Author Contributions

J.Z. led computational pipeline development, data analysis, visualization, and manuscript drafting. E.A.C. and R.D.W. designed and performed large-scale microscopy experiments and contributed datasets, with additional validation data from L.J.S. J.Z., E.A.C., P.C., and T.T.K. curated and annotated datasets. J.Z., A.B.P., and O.A.I. designed the deep learning framework, performed model training, and analyzed human evaluation experiments. P.C. and P.A.M. assisted in experimental design. J.Z., P.C., R.D.W., and O.A.I. wrote and revised the manuscript. A.B.P., R.D.W., and O.A.I. supervised the work. All authors discussed the results and approved the final manuscript.

## Data Availability

The training and evaluation datasets generated and analyzed during this study have been deposited in the BioImage Archive under accession code S-BIAD2328 and are publicly available at https://doi.org/10.6019/S-BIAD2328. The source code is publicly available on GitHub at https://github.com/IgoshinLab/StyleAutoEncoder.

## Supplementary Information

### Stratified Sampling

Movies in our dataset begin without aggregates, and visible aggregates typically form after more than 6 hours of observation. As a result, early frames across strains appear phenotypically uniform, while phenotypic diversity becomes more apparent at later time points. Additionally, mutations that impair sporulation result in no aggregate formation throughout the movie. Consequently, frames without aggregates substantially outnumber those with visible aggregates. Moreover, phenotypic categories differ in their number of experimental replicates and in the temporal duration of aggregate visibility, leading to substantial class imbalance in randomly sampled training batches (Figure S1A).

To address this imbalance, we employed stratified sampling based on rough phenotype labels generated from a preliminary classification model. We trained an Inception-V3 classifier on selected movie frames: frame 1 (consistently labeled as “No aggregate”) and frames 1081, 1241, and 1441 (when available), which were manually labeled by human annotators into 13 broad phenotype categories. To improve generalization and mitigate overfitting in underrepresented classes, we applied data augmentation via random rotations and cropping. Despite inherent ambiguity between some categories, we partitioned the labeled data into training and validation sets using a 4:1 ratio to preserve category balance.

For training, one movie from each of 200 distinct strains was reserved for validation, with the remaining movies used for training. The model was trained using cross-entropy loss, monitored with a rough-label-balanced cross-entropy metric for early stopping (batch size = 600, evaluation every 10 epochs, patience = 7 epochs). Training converged after approximately 650 epochs (Figure S1B). Although the validation accuracy plateaued at 50% due to label ambiguity, this was sufficient for generating preliminary (“rough”) labels (Figure S1C). We then applied the trained classifier to assign rough labels to the full dataset, enabling stratified sampling during training. This ensured that each batch contained more balanced representation across the rough phenotype categories (Figure S1D).

### Hyper-parameter Search

Our goal is to ensure the model encodes only phenotypic information into the feature space, disentangling it from nuisance variables, particularly positional cues. Additionally, for analytical simplicity, we aim to determine the minimal number of encoding dimensions necessary to achieve effective disentanglement. Initially, training the model from scratch on our dataset led to entanglement of positional information with phenotype. Specifically, altering the input image (Figure S2A) changed both the reconstructed phenotype and the positions of aggregates (columns 1/2 vs. columns 3/4), which contradicts our objective since phenotype should be independent of aggregate positioning. Furthermore, changes in the noise vector alone had no impact on either phenotype or aggregate positions (Figure S2H), underscoring this entanglement.

To overcome this issue, we implemented a two-step training approach. The model was first pre-trained on Shimkets’ dataset, benefiting from its greater variability in aggregate positioning, thus facilitating the separation of positional information from phenotypic traits. Subsequently, we fine-tuned this pre-trained model on our dataset. This strategy successfully achieved disentanglement, as illustrated in Figure S2I: changing the input strain from DK1622 to DK2232 altered the reconstructed phenotype, while varying the noise vector distinctly shifted aggregate positions.

Following this successful disentanglement, we conducted a hyper-parameter search to identify the minimal encoding dimensionality, varying it from 10 to 15 dimensions. Models trained on Shimkets’ dataset (252k images) revealed clear trends (Figures S2B–G): For encoding dimensions between 10 and 12, changes in input images or noise vectors had minimal effects on reconstructed phenotypes and aggregate positions (Figures S2B–D). However, at higher dimensionalities (13 to 15), the noise vector distinctly influenced aggregate positions (Figures S2E–G). Importantly, only at an encoding dimension of 13 did changing the input image from DK1622 to DK2232 consistently result in distinct reconstructed phenotypes; this effect diminished at encoding dimensions of 14 and 15. Therefore, given our architecture and training parameters, an encoding dimension of 13 appears optimal, effectively capturing phenotype information while clearly disentangling positional information.

### Human Evaluation

To further investigate the human evaluation results, we conducted separate analyses for the non-expert and expert participant groups. In the analysis of the 21 non-expert participants, fidelity ratings were consistent with the combined results, showing no significant difference between real images (mean ± SD: 3.34±1.19), our model’s images (3.52±1.12), and StyleGAN2 images (3.36±1.18). In contrast, Style-GAN3 images received significantly lower scores (1.08±0.35)(Figure S3A, Table 5). In the alignment task, non-experts ranked the right crop highest (4.16±0.95), followed by our model (3.61±1.16), the experimental replicate (2.97±1.33), and StyleGAN2 (2.77±1.19)(Figure S3B). Our model’s reconstructions were rated significantly higher than both StyleGAN2 (p¡0.001, g=1.19) and the experimental replicate (p¡0.001, g=0.79). For this group, the difference between StyleGAN2 and the experimental replicate was not statistically significant (p=0.35, g=0.29)(Table 6).

The analysis of the 18 expert participants revealed a more nuanced assessment of image fidelity. Experts rated images from our model as having the highest fidelity (3.38±1.33), a statistically significant, albeit small, difference compared to both real images (3.18±1.40; p=0.0068, g=0.14) and StyleGAN2 images (3.22±1.38; p=0.0481, g=0.12). The difference between real and StyleGAN2 images was not significant (p=0.92, g=-0.03). As with other groups, StyleGAN3 images were rated significantly lower than all other categories (1.12±0.55; all p¡0.001)(Figure S3C, Table 5). For the alignment evaluation, the mean scores from experts followed the same hierarchical pattern observed in the combined analysis: the right crop received the highest score (3.97±1.14), followed by our model (3.51±1.17), the experimental replicate (2.92±1.30), and the StyleGAN2 reconstruction (2.69±1.24)(Figure S3D, Table 6). This indicates that experts found our model’s reconstructions to be a significantly better match to the reference phenotype than both experimental replicates and StyleGAN2 reconstructions.

### Case Studies for Feature Evaluation

To provide a qualitative understanding of the performance differences between model-extracted and human-defined features, we selected representative examples from our dataset for two key scenarios.

First, we identified cases where our model’s features succeeded while human-defined features failed (Figure S4A-C). In these examples, which include strains like DK4492, Ω4473, and DK101, the human-defined features reported a strong negative correlation between paired image crops. In stark contrast, our model produced high cosine similarity scores for these same pairs, correctly identifying their shared phenotypic identity. These cases typically occur in images with sparse or non-obvious aggregates, where the model’s ability to capture subtle textural and spatial patterns excels.

Second, as a positive control, we identified cases where both methods performed well (Figure S4D-F). In strains like DK1218 and DK1410, both the human-defined and model-extracted features yielded high cosine similarity scores, confirming that both methods agree when presented with clear, unambiguous phenotypes. Notably, even in these cases, the model’s similarity scores were consistently high, demonstrating its reliability.

Finally, we found no instances where human-defined features succeeded and the model failed. This highlights the robustness and superior coverage of the model-derived feature space, which appears to encapsulate the information from traditional metrics while adding a deeper layer of quantitative description.

### Loss Functions

**Discriminator Loss** The discriminator loss is decomposed into four sub-losses:

1. Real Pair Loss:

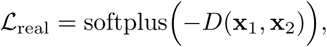

which encourages high similarity between crops from the same image.

2. Fake Pair Loss:

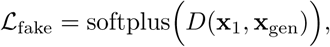

which penalizes similarity when one image is generated.

3. R1 Regularization:

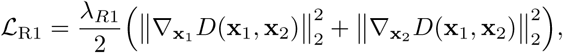

which stabilizes training by penalizing the squared *L*_2_-norm of the gradients, with *λ_R_*_1_ controlling the regularization strength.

4. **Gradient Penalty (WGAN-GP):** By interpolating between **x**_1_ and **x**_gen_, we define

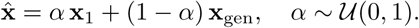

Then, the gradient penalty is given by

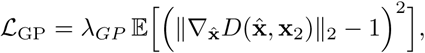

where *λ_GP_*is a weighting hyperparameter that enforces Lipschitz continuity.

The total discriminator loss is thus:

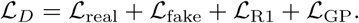

**Encoder-generator Loss** The encoder–generator branch is trained to minimize the following losses:

1. KL-Divergence Loss:

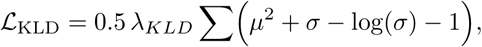

which regularizes the latent space to follow a unit Gaussian distribution. This ensures that the latent representations are well-behaved and facilitates effective interpolation.

2. Adversarial Loss:

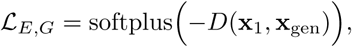

ensuring that generated images effectively “deceive” the discriminator.

3. Path Length Penalty:

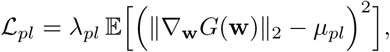

which regularizes the generator’s mapping by enforcing a consistent perceptual change. In our implementation, the gradient ∇**_w_***G*(**w**) is scaled by a noise term (pl noise) to account for the spatial dimensions of the generated image; this detail is inherent in the StyleGAN2 design and is omitted here for brevity.

### Classification and Performance Evaluation

#### Support Vector Machine (SVM) Classification

To classify phenotypes, we employed a Support Vector Machine (SVM) classifier with a radial basis function (RBF) kernel. The classification was performed on the 13-dimensional feature vectors extracted from images at specific time points (0 h and 24 h).

#### Cross-Validation

We used a stratified 3-fold cross-validation scheme to train and evaluate the SVM models, ensuring that each fold contained a proportional representation of each class. For the analysis of motility mutants, which involved multiple movies from the same strain, we utilized StratifiedGroupKFold. This approach guarantees that all data from a single strain (genotype) were kept within the same fold during splitting, thus preventing data leakage between the training and testing sets and providing a more robust estimate of generalization performance.

#### Performance Metrics

The model’s ability to distinguish between classes was quantified using the Area Under the Receiver Operating Characteristic (ROC-AUC) curve. The reported AUC is the mean score across the 3 cross-validation folds. The 95% confidence interval (CI) was calculated as 1.96×SEM, where SEM is the standard error of the mean of the AUC scores from the folds.

#### Dimensionality Reduction and Visualization

For visualization, we used Principal Component Analysis (PCA) to reduce the 13-dimensional feature space to its first two principal components. The PCA was fitted on the combined feature data from both classes at a given time point. To visualize the SVM decision boundary, we trained an SVM on the 2D PCA-transformed data. We then created a fine grid over the 2D space and used the trained model to predict the class probability for each point. These probabilities are displayed as contour plots overlaid on the PCA scatter plot, illustrating the separation between classes.

**Figure S1:**
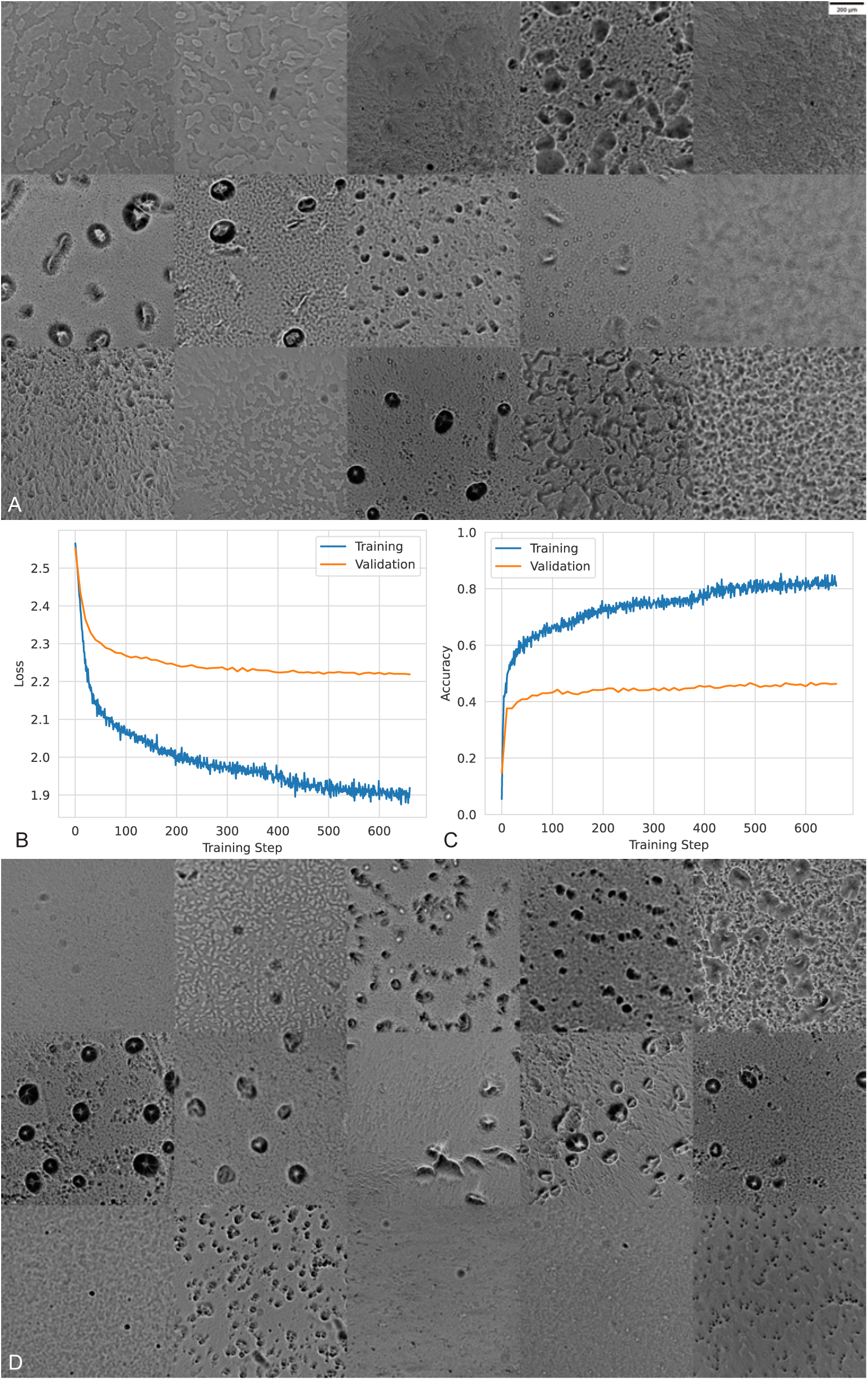
Stratified Sampling. (A) A training batch constructed via fully random sampling, illustrating overrepresentation of early-stage, non-aggregated frames. (B) Training and validation loss curves of the rough-label classifier. (C) Accuracy curves during training, highlighting label ambiguity. (D) A training batch constructed via stratified sampling, demonstrating improved coverage across rough phenotype categories.

**Table 1:** Summary of *Myxococcus xanthus* strains and corresponding swarm motility data. The table lists the genetically distinct strains included in the time-lapse microscopy dataset. For each strain, the total number of replicate movies is provided, along with full bibliographic citations.

| Strain | Number of Movies | motility | Bib |
| --- | --- | --- | --- |
| DK101 | 9 | A+S+ | [19, 65] |
| DK1622 | 13 | A+S+ | [62] |
| DK3119 | 3 | A-S- | [67] |
| DK4160 | 3 | A-S- | [106] |
| DK2160 | 3 | A-S- | [50] |
| DK1259 | 2 | A-S- | [19, 80] |
| DK1409 | 3 | A-S- | [62] |
| DK4050 | 2 | A-S- | [115] |
| DK4153 | 3 | A-S- | [115, 114] |
| DK2618 | 3 | A-S+ | [112] |
| DK2616 | 3 | A-S+ | [110] |
| DK2614 | 2 | A-S+ | [67] |
| DK2608 | 3 | A-S+ | [67] |
| DK2232 | 3 | A-S+ | [69] |
| DK1213 | 3 | A-S+ | [19] |
| DK1215 | 3 | A-S+ | [19] |
| DK1217 | 3 | A-S+ | [19] |
| DK1410 | 3 | A-S+ | [62] |
| DK7679 | 3 | A-S+ | [62] |
| DK1227 | 3 | A-S+ | [19] |
| DK1224 | 3 | A-S+ | [62] |
| DK1221 | 3 | A-S+ | [19] |
| DK1220 | 3 | A-S+ | [19] |
| DK1218 | 3 | A-S+ | [19] |
| DK3475 | 3 | A+S- | [67] |
| DK8615 | 3 | A+S- | [122] |
| DK3481 | 3 | A+S- | [110] |
| DK7558 | 2 | A+S- | [85] |
| DK2158 | 3 | A+S- | [50] |
| DK3186 | 3 | A+S- | [64] |
| DK3068 | 3 | A+S- | [31] |
| DK2112 | 2 | A+S- | [50] |
| DK1932 | 2 | A+S- | [105] |
| DK1680 | 3 | A+S- | [57, 121] |
| DK1670 | 3 | A+S- | [50] |
| DK1300 | 6 | A+S- | [116] |
| DK1253 | 6 | A+S- | [80] |
| DK8620 | 3 | A+S- | [62] |
| ASX1 | 3 | - | [62] |
| DK1013 | 2 | - | [110] |
| DK1016 | 3 | - | [111] |

Table 1: – continued from previous page
| Strain | Number of Movies | motility | Bib |
| --- | --- | --- | --- |
| DK1031 | 2 | - | [110] |
| DK10524 | 3 | - | [97] |
| DK10527 | 3 | - | [97] |
| DK10536 | 3 | - | [62] |
| DK10546 | 3 | - | [2] |
| DK10603 | 1 | - | [126] |
| DK10604 | 3 | - | [73] |
| DK1061 | 2 | - | [62] |
| DK11212 | 3 | - | [97] |
| DK119 | 3 | - | [62] |
| DK121 | 2 | - | [40] |
| DK129 | 3 | - | [62] |
| DK149 | 3 | - | [98] |
| DK1525 | 3 | - | [112] |
| DK1529 | 3 | - | [95] |
| DK1627 | 2 | - | [120] |
| DK1648 | 3 | - | [62] |
| DK1666 | 3 | - | [50] |
| DK1669 | 3 | - | [62] |
| DK2109 | 2 | - | [119] |
| DK2110 | 2 | - | [62] |
| DK2118 | 3 | - | [62] |
| DK2134 | 3 | - | [62] |
| DK2138 | 3 | - | [62] |
| DK2143 | 1 | - | [124] |
| DK2161 | 1 | - | [50] |
| DK2210 | 2 | - | [62] |
| DK2213 | 2 | - | [64] |
| DK2231 | 2 | - | [103] |
| DK233 | 3 | - | [62] |
| DK2737 | 1 | - | [62] |
| DK315 | 3 | - | [77] |
| DK3151 | 3 | - | [64] |
| DK3250 | 3 | - | [62] |
| DK3252 | 3 | - | [31] |
| DK3258 | 2 | - | [62] |
| DK3261 | 3 | - | [31] |
| DK3354 | 2 | - | [62] |
| DK3430 | 3 | - | [62] |
| DK3468 | 2 | - | [94] |
| DK3516 | 3 | - | [62] |
| DK3517 | 3 | - | [63] |

Table 1: – continued from previous page
| Strain | Number of Movies | motility | Bib |
| --- | --- | --- | --- |
| DK3517 | 3 | - | [63] |
| DK3518 | 3 | - | [62] |
| DK3551 | 3 | - | [63] |
| DK3619 | 3 | - | [115] |
| DK3955 | 5 | - | [107] |
| DK3959 | 2 | - | [62] |
| DK406 | 3 | - | [102] |
| DK412 | 2 | - | [31] |
| DK4141 | 3 | - | [88] |
| DK4162 | 3 | - | [106] |
| DK418 | 3 | - | [31] |
| DK425 | 3 | - | [31] |
| DK426 | 3 | - | [31] |
| DK4285 | 3 | - | [94] |
| DK429 | 3 | - | [31] |
| DK4290 | 3 | - | [89] |
| DK4292 | 3 | - | [89] |
| DK4293 | 6 | - | [120] |
| DK4296 | 3 | - | [89] |
| DK4297 | 3 | - | [28] |
| DK4298 | 3 | - | [62] |
| DK4299 | 3 | - | [81] |
| DK4300 | 5 | - | [104] |
| DK4302 | 3 | - | [94] |
| DK4308 | 3 | - | [94] |
| DK4309 | 2 | - | [117] |
| DK4310 | 3 | - | [59] |
| DK4312 | 3 | - | [94] |
| DK4313 | 2 | - | [94] |
| DK4316 | 3 | - | [62] |
| DK4322 | 6 | - | [94] |
| DK4323 | 6 | - | [94] |
| DK4324 | 6 | - | [94] |
| DK4325 | 3 | - | [94] |
| DK4366 | 3 | - | [94] |
| DK4371 | 3 | - | [59] |
| DK4372 | 3 | - | [94] |
| DK4375 | 5 | - | [94] |
| DK4376 | 3 | - | [94] |
| DK4377 | 3 | - | [94] |
| DK4379 | 3 | - | [59] |
| DK4380 | 3 | - | [94] |

Table 1: – continued from previous page
| Strain | Number of Movies | motility | Bib |
| --- | --- | --- | --- |
| DK4386 | 6 | - | [59, 94] |
| DK4387 | 3 | - | [94] |
| DK4389 | 3 | - | [94] |
| DK4396 | 3 | - | [31] |
| DK4399 | 6 | - | [94] |
| DK440 | 3 | - | [95] |
| DK4401 | 3 | - | [94] |
| DK4408 | 3 | - | [94] |
| DK4414 | 3 | - | [120] |
| DK4435 | 3 | - | [96] |
| DK4442 | 3 | - | [94] |
| DK4445 | 3 | - | [62] |
| DK4451 | 3 | - | [62] |
| DK4459 | 3 | - | [64] |
| DK4464 | 6 | - | [62] |
| DK4474 | 3 | - | [89] |
| DK4481 | 6 | - | [28] |
| DK4492 | 3 | - | [89] |
| DK4499 | 3 | - | [68] |
| DK4500 | 3 | - | [28] |
| DK4506 | 3 | - | [89, 28] |
| DK4517 | 6 | - | [28] |
| DK4519 | 6 | - | [84] |
| DK4521 | 12 | - | [89, 85] |
| DK4529 | 3 | - | [89] |
| DK4530 | 2 | - | [89] |
| DK4535 | 3 | - | [62] |
| DK4564 | 3 | - | [31] |
| DK4566 | 3 | - | [31] |
| DK4587 | 2 | - | [92] |
| DK4619 | 3 | - | [101] |
| DK4678 | 3 | - | [101] |
| DK4696 | 3 | - | [62] |
| DK473 | 6 | - | [95] |
| DK4746 | 6 | - | [82] |
| DK4750 | 3 | - | [62] |
| DK476 | 3 | - | [94] |
| DK4764 | 5 | - | [101] |
| DK4765 | 3 | - | [101] |
| DK4778 | 3 | - | [101] |
| DK48 | 3 | - | [62] |
| DK4861 | 8 | - | [31] |

Table 1: – continued from previous page
| Strain | Number of Movies | motility | Bib |
| --- | --- | --- | --- |
| DK4863 | 3 | - | [31] |
| DK4870 | 3 | - | [31] |
| DK495 | 3 | - | [75] |
| DK5055 | 1 | - | [31] |
| DK5056 | 3 | - | [31] |
| DK5057 | 3 | - | [31] |
| DK50576 | 2 | - | [62] |
| DK5058 | 5 | - | [31] |
| DK5061 | 3 | - | [31] |
| DK5063 | 3 | - | [31] |
| DK5066 | 3 | - | [31] |
| DK5067 | 3 | - | [31] |
| DK5068 | 3 | - | [31] |
| DK5069 | 2 | - | [31] |
| DK5070 | 3 | - | [92] |
| DK5073 | 3 | - | [31] |
| DK5074 | 3 | - | [31] |
| DK5090 | 2 | - | [31] |
| DK510 | 3 | - | [19] |
| DK516 | 3 | - | [31] |
| DK5200 | 5 | - | [89] |
| DK5203 | 3 | - | [92] |
| DK5204 | 3 | - | [96] |
| DK5205 | 3 | - | [82] |
| DK5206 | 9 | - | [89] |
| DK5208 | 6 | - | [97] |
| DK5209 | 8 | - | [83] |
| DK5216 | 3 | - | [89] |
| DK5217 | 3 | - | [89] |
| DK5225 | 3 | - | [89] |
| DK5246 | 5 | - | [89] |
| DK5247 | 3 | - | [89] |
| DK5251 | 3 | - | [89] |
| DK5257 | 3 | - | [89] |
| DK5270 | 2 | - | [89] |
| DK5274 | 3 | - | [91] |
| DK5275 | 3 | - | [91] |
| DK5276 | 2 | - | [91] |
| DK5277 | 3 | - | [91] |
| DK5279 | 5 | - | [87] |
| DK5280 | 3 | - | [85] |
| DK5281 | 3 | - | [62] |

Table 1: – continued from previous page
| Strain | Number of Movies | motility | Bib |
| --- | --- | --- | --- |
| DK5285 | 3 | - | [89] |
| DK5510 | 3 | - | [62] |
| DK5511 | 2 | - | [67] |
| DK5870 | 2 | - | [62] |
| DK590 | 6 | - | [99] |
| DK6000 | 2 | - | [100] |
| DK6001 | 3 | - | [62] |
| DK6058 | 2 | - | [62] |
| DK6206 | 3 | - | [76] |
| DK653 | 3 | - | [75] |
| DK6601 | 2 | - | [82] |
| DK6607 | 3 | - | [82] |
| DK6620 | 3 | - | [74] |
| DK6621 | 3 | - | [74] |
| DK6643 | 3 | - | [82] |
| DK6665 | 3 | - | [2] |
| DK6759 | 3 | - | [62] |
| DK68 | 3 | - | [62] |
| DK7052 | 3 | - | [62] |
| DK7053 | 3 | - | [62] |
| DK7056 | 8 | - | [62] |
| DK7057 | 6 | - | [62] |
| DK7058 | 3 | - | [62] |
| DK7088 | 2 | - | [67] |
| DK7090 | 5 | - | [62] |
| DK7091 | 6 | - | [62] |
| DK7154 | 3 | - | [62] |
| DK7160 | 3 | - | [71] |
| DK718 | 3 | - | [75] |
| DK7293 | 2 | - | [62] |
| DK7294 | 3 | - | [62] |
| DK7295 | 3 | - | [62] |
| DK731 | 3 | - | [75] |
| DK739 | 3 | - | [75] |
| DK741 | 2 | - | [112] |
| DK7507 | 3 | - | [62] |
| DK7509 | 3 | - | [62] |
| DK7512 | 3 | - | [62] |
| DK7517 | 3 | - | [2] |
| DK752 | 3 | - | [31] |
| DK7523 | 3 | - | [62] |
| DK7559 | 2 | - | [62] |

Table 1: – continued from previous page
| Strain | Number of Movies | motility | Bib |
| --- | --- | --- | --- |
| DK756 | 2 | - | [75] |
| DK7563 | 3 | - | [85] |
| DK7575 | 3 | - | [62] |
| DK7578 | 3 | - | [62] |
| DK7585 | 2 | - | [62] |
| DK762 | 3 | - | [113] |
| DK7627 | 2 | - | [62] |
| DK7636 | 3 | - | [85] |
| DK7639 | 1 | - | [62] |
| DK7646 | 2 | - | [85] |
| DK7669 | 3 | - | [62] |
| DK7761 | 3 | - | [62] |
| DK7763 | 3 | - | [62] |
| DK777 | 3 | - | [101] |
| DK7824 | 2 | - | [72] |
| DK7825 | 3 | - | [60] |
| DK7826 | 3 | - | [72] |
| DK7827 | 2 | - | [71] |
| DK785 | 3 | - | [62] |
| DK7862 | 2 | - | [62] |
| DK8617 | 3 | - | [120] |
| DK8624 | 3 | - | [62] |
| DK9521 | 3 | - | [84] |
| FruATc | 3 | - | [62] |
| MXAN1010 | 3 | - | [66] |
| MXAN1909 | 2 | - | [61] |
| MXAN3442 | 3 | - | [70] |
| MXAN3474 | 2 | - | [127] |
| MXAN3553 | 2 | - | [86] |
| MXAN3557 | 3 | - | [86] |
| MXAN4251 | 3 | - | [108] |
| MXAN4398 | 6 | - | [118] |
| MXAN4400 | 3 | - | [62] |
| MXAN4494 | 1 | - | [79] |
| MXAN7059 | 6 | - | [123] |
| MXAN7089 | 6 | - | [58] |
| MXAN7164 | 3 | - | [61] |
| $\Omega 4531$ | 1 | - | [28] |
| esgWen | 2 | - | [62] |
| $\Omega 4469$ | 3 | - | [94] |
| $\Omega 4473$ | 3 | - | [94] |
| **Total** | 937 |  |  |

**Table 2:**
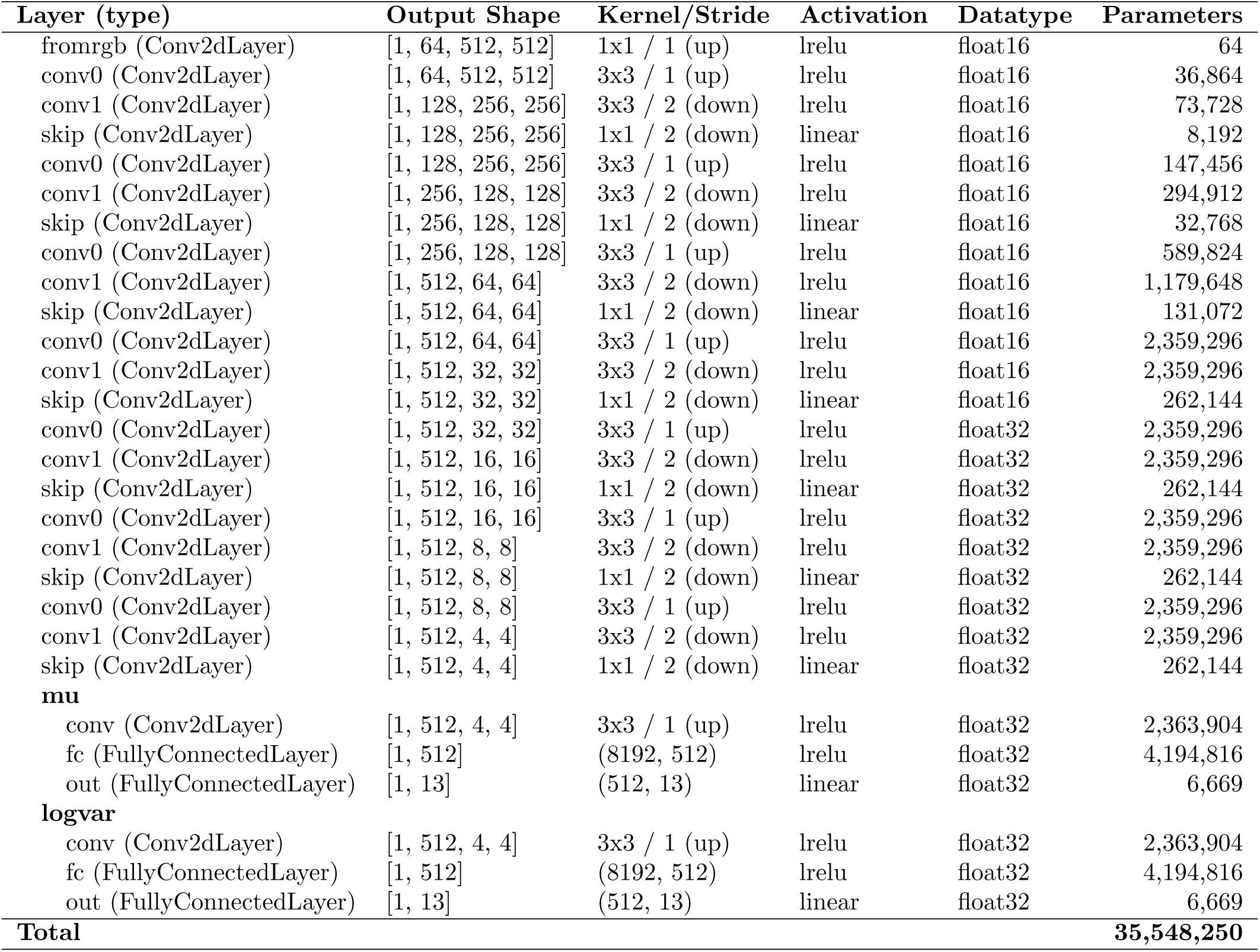
Detailed architecture of the Encoder (E) network.

| Layer (type) | Output Shape | Kernel/Stride | Activation | Datatype | Parameters |
| --- | --- | --- | --- | --- | --- |
| fromrgb (Conv2dLayer) | [1, 64, 512, 512] | 1x1 / 1 (up) | lrelu | float16 | 64 |
| conv0 (Conv2dLayer) | [1, 64, 512, 512] | 3x3 / 1 (up) | lrelu | float16 | 36,864 |
| conv1 (Conv2dLayer) | [1, 128, 256, 256] | 3x3 / 2 (down) | lrelu | float16 | 73,728 |
| skip (Conv2dLayer) | [1, 128, 256, 256] | 1x1 / 2 (down) | linear | float16 | 8,192 |
| conv0 (Conv2dLayer) | [1, 128, 256, 256] | 3x3 / 1 (up) | lrelu | float16 | 147,456 |
| conv1 (Conv2dLayer) | [1, 256, 128, 128] | 3x3 / 2 (down) | lrelu | float16 | 294,912 |
| skip (Conv2dLayer) | [1, 256, 128, 128] | 1x1 / 2 (down) | linear | float16 | 32,768 |
| conv0 (Conv2dLayer) | [1, 256, 128, 128] | 3x3 / 1 (up) | lrelu | float16 | 589,824 |
| conv1 (Conv2dLayer) | [1, 512, 64, 64] | 3x3 / 2 (down) | lrelu | float16 | 1,179,648 |
| skip (Conv2dLayer) | [1, 512, 64, 64] | 1x1 / 2 (down) | linear | float16 | 131,072 |
| conv0 (Conv2dLayer) | [1, 512, 64, 64] | 3x3 / 1 (up) | lrelu | float16 | 2,359,296 |
| conv1 (Conv2dLayer) | [1, 512, 32, 32] | 3x3 / 2 (down) | lrelu | float16 | 2,359,296 |
| skip (Conv2dLayer) | [1, 512, 32, 32] | 1x1 / 2 (down) | linear | float16 | 262,144 |
| conv0 (Conv2dLayer) | [1, 512, 32, 32] | 3x3 / 1 (up) | lrelu | float32 | 2,359,296 |
| conv1 (Conv2dLayer) | [1, 512, 16, 16] | 3x3 / 2 (down) | lrelu | float32 | 2,359,296 |
| skip (Conv2dLayer) | [1, 512, 16, 16] | 1x1 / 2 (down) | linear | float32 | 262,144 |
| conv0 (Conv2dLayer) | [1, 512, 16, 16] | 3x3 / 1 (up) | lrelu | float32 | 2,359,296 |
| conv1 (Conv2dLayer) | [1, 512, 8, 8] | 3x3 / 2 (down) | lrelu | float32 | 2,359,296 |
| skip (Conv2dLayer) | [1, 512, 8, 8] | 1x1 / 2 (down) | linear | float32 | 262,144 |
| conv0 (Conv2dLayer) | [1, 512, 8, 8] | 3x3 / 1 (up) | lrelu | float32 | 2,359,296 |
| conv1 (Conv2dLayer) | [1, 512, 4, 4] | 3x3 / 2 (down) | lrelu | float32 | 2,359,296 |
| skip (Conv2dLayer) | [1, 512, 4, 4] | 1x1 / 2 (down) | linear | float32 | 262,144 |
| **mu** |  |  |  |  |  |
| conv (Conv2dLayer) | [1, 512, 4, 4] | 3x3 / 1 (up) | lrelu | float32 | 2,363,904 |
| fc (FullyConnectedLayer) | [1, 512] | (8192, 512) | lrelu | float32 | 4,194,816 |
| out (FullyConnectedLayer) | [1, 13] | (512, 13) | linear | float32 | 6,669 |
| **logvar** |  |  |  |  |  |
| conv (Conv2dLayer) | [1, 512, 4, 4] | 3x3 / 1 (up) | lrelu | float32 | 2,363,904 |
| fc (FullyConnectedLayer) | [1, 512] | (8192, 512) | lrelu | float32 | 4,194,816 |
| out (FullyConnectedLayer) | [1, 13] | (512, 13) | linear | float32 | 6,669 |
| **Total** |  |  |  |  | **35,548,250** |

**Figure S2:**
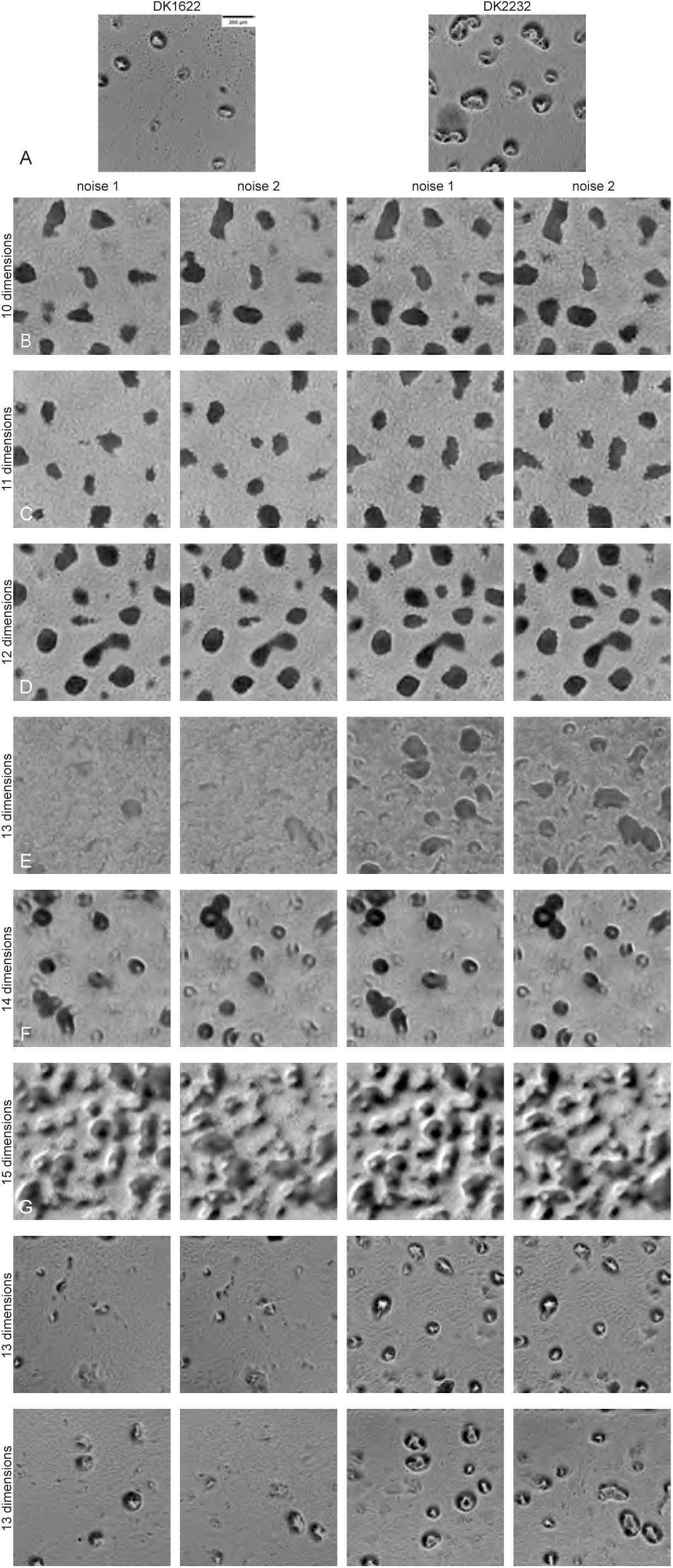
Hyper-parameter search for encoding dimensions. (A) Reference input images. (B-G) Reconstructed images from models trained on Shimkets’ dataset (252 × 10^3^ images) with random noise and encoding dimensions ranging from 10 to 15. Column 1 and 3 use the same noise, and column 2 and 4 use another noise. (H) Reconstructed images from the model trained on Welch’s dataset (1, 512 × 10^3^ images) using random noise and an encoding dimension of 13. (I) Reconstructed images from the model first pre-trained on Shimkets’ dataset (302 × 10^3^ images), then fine-tuned on our dataset (1, 512 × 10^3^ images) with random noise and an encoding dimension of 13. Column 1 and 3 use the same noise, and column 2 and 4 use another noise.

**Figure S3:**
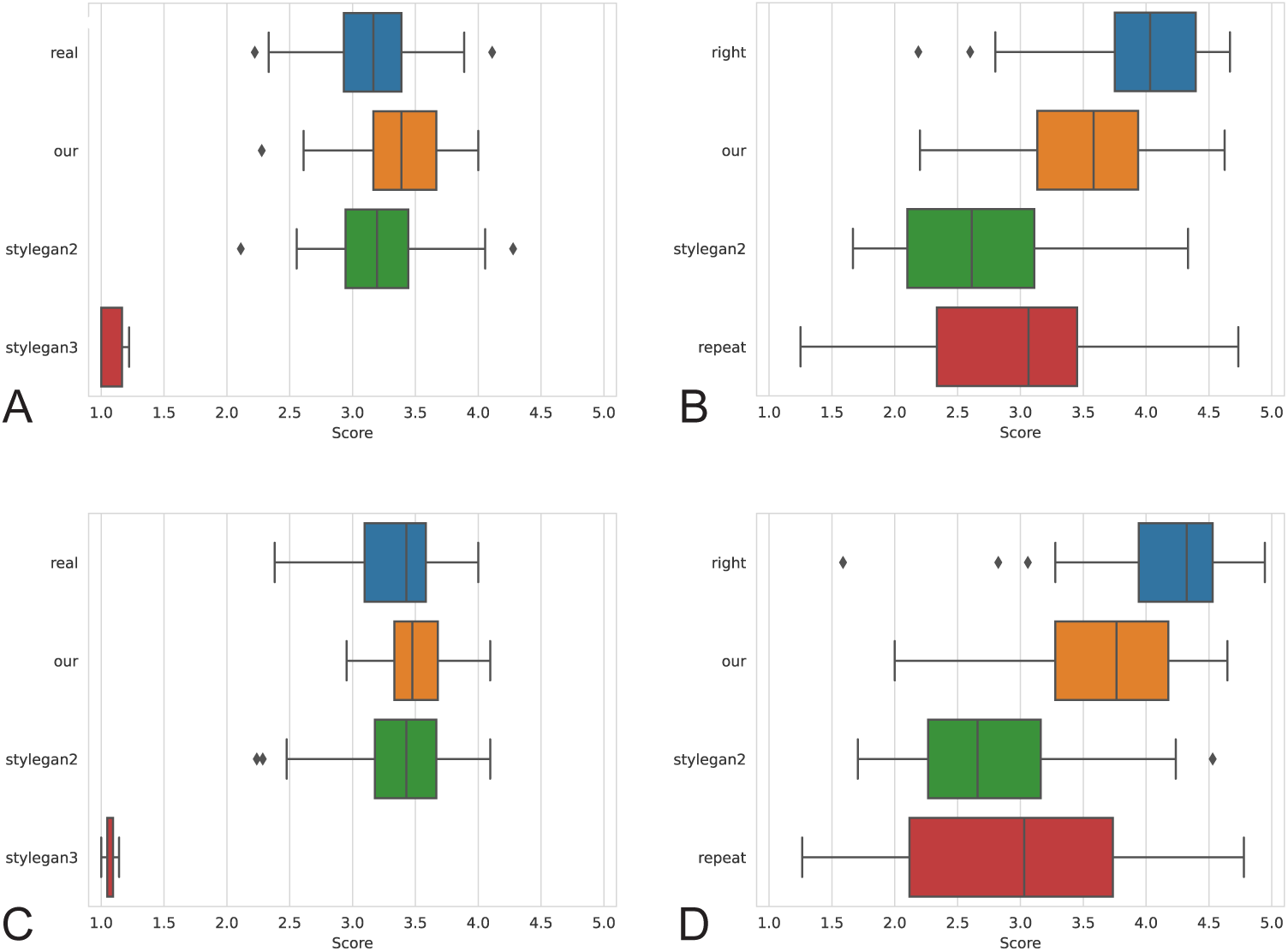
Human evaluation of fidelity and alignment across expert and non-expert groups. (A, C) Distribution of fidelity ratings from (A) experts and (C) non-experts, comparing real images, our model, StyleGAN2, and StyleGAN3-generated images. (B, D) Distribution of alignment ratings from (B) experts and (D) non-experts, comparing reference images to right crops, experimental replicates, and reconstructions from our model and StyleGAN2.

**Figure S4:**
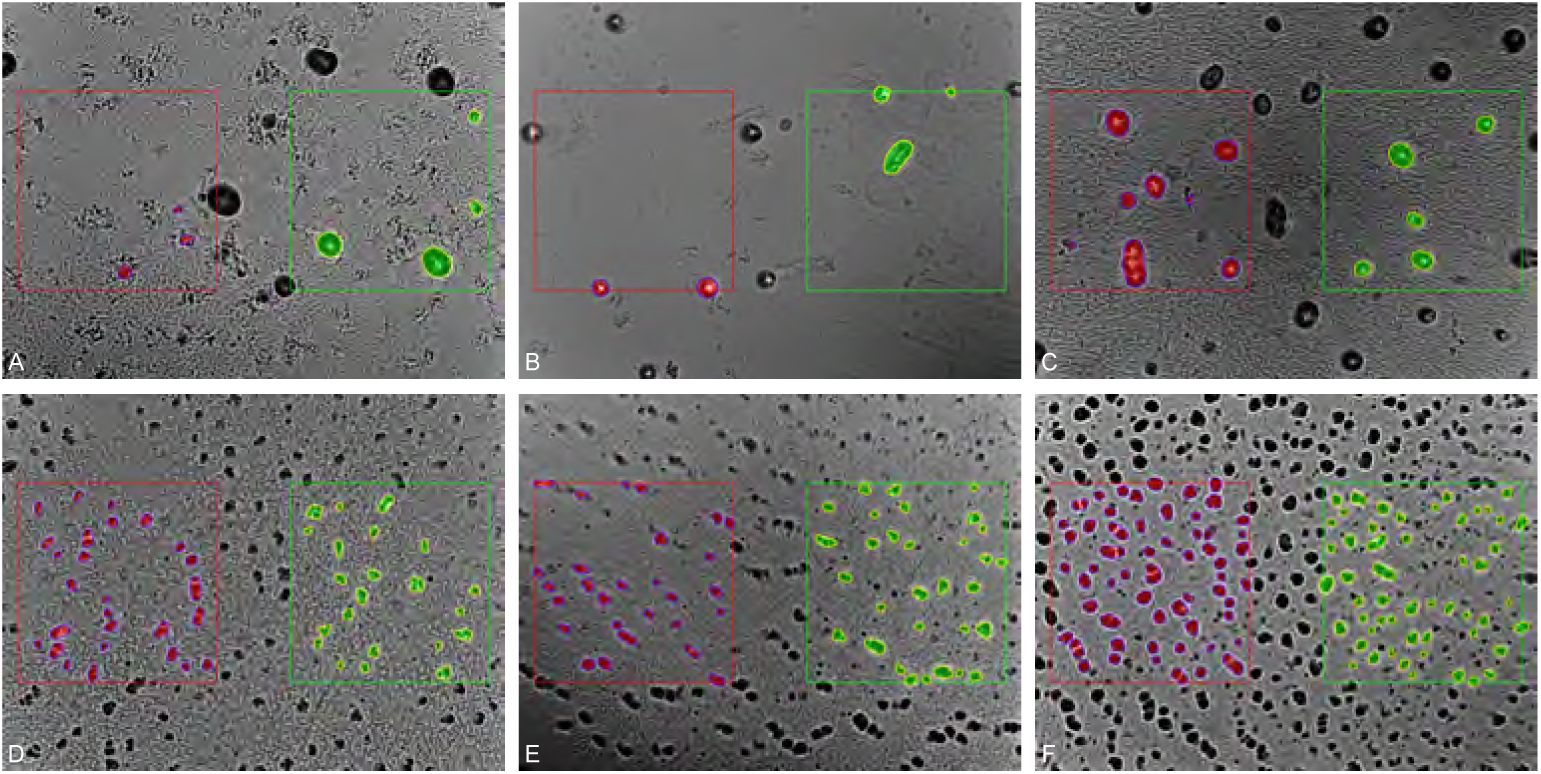
Case study examples illustrating the performance of model-extracted vs. human-defined features. (A-C) Three examples where the model succeeded while human features failed, with examples from strains (A) DK4492 (model sim: 0.88, human sim: −0.75), (B) Ω4473 (model sim: 0.85, human sim: −0.68), and (C) DK101 (model sim: 0.84, human sim: −0.51). (D-F) Three examples where both methods succeeded, featuring strains (D) DK1218 (model sim: 0.99, human sim: 0.97), and two examples from DK1410 ((E) model sim: 0.99, human sim: 0.97; and (F) model sim: 0.98, human sim: 0.98).

**Figure S5:**
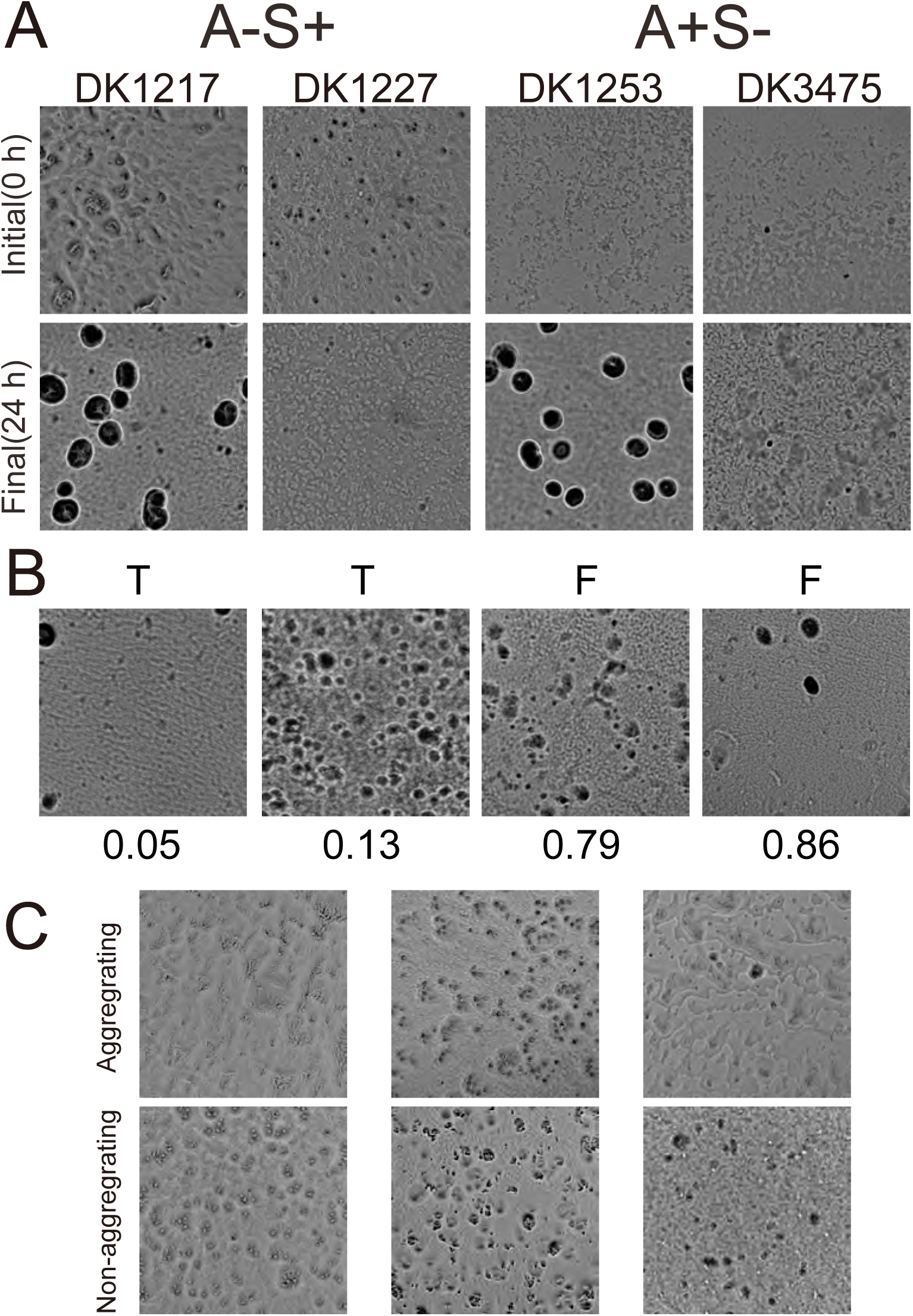
(A) Initial (0 h) and final (24 h) images of motility mutant strains A-S+ (DK1217 and DK1227) and A+S-(DK1253 and DK3475). Both motility mutants can lead to aggregates at 24 h (DK1217 and DK1253) or fail to form aggregates at 24 h (DK1227 and DK3475). (B) Misclassified examples from the SVM: two false negatives (true aggregating, low score) and two false positives (true non-aggregating, high score). Top: human labels; bottom: model-predicted aggregation scores (1 = aggregating, 0 = non-aggregating). (C) Samples of initial (0 h) images of different WT experiments in similar conditions grouped by their ability to form aggregates at the 24 h time point (see Figure 3C for the final time frame of the same experiments) with the top row showing aggregating and the bottom row non-aggregating experiments.

**Table 3:**
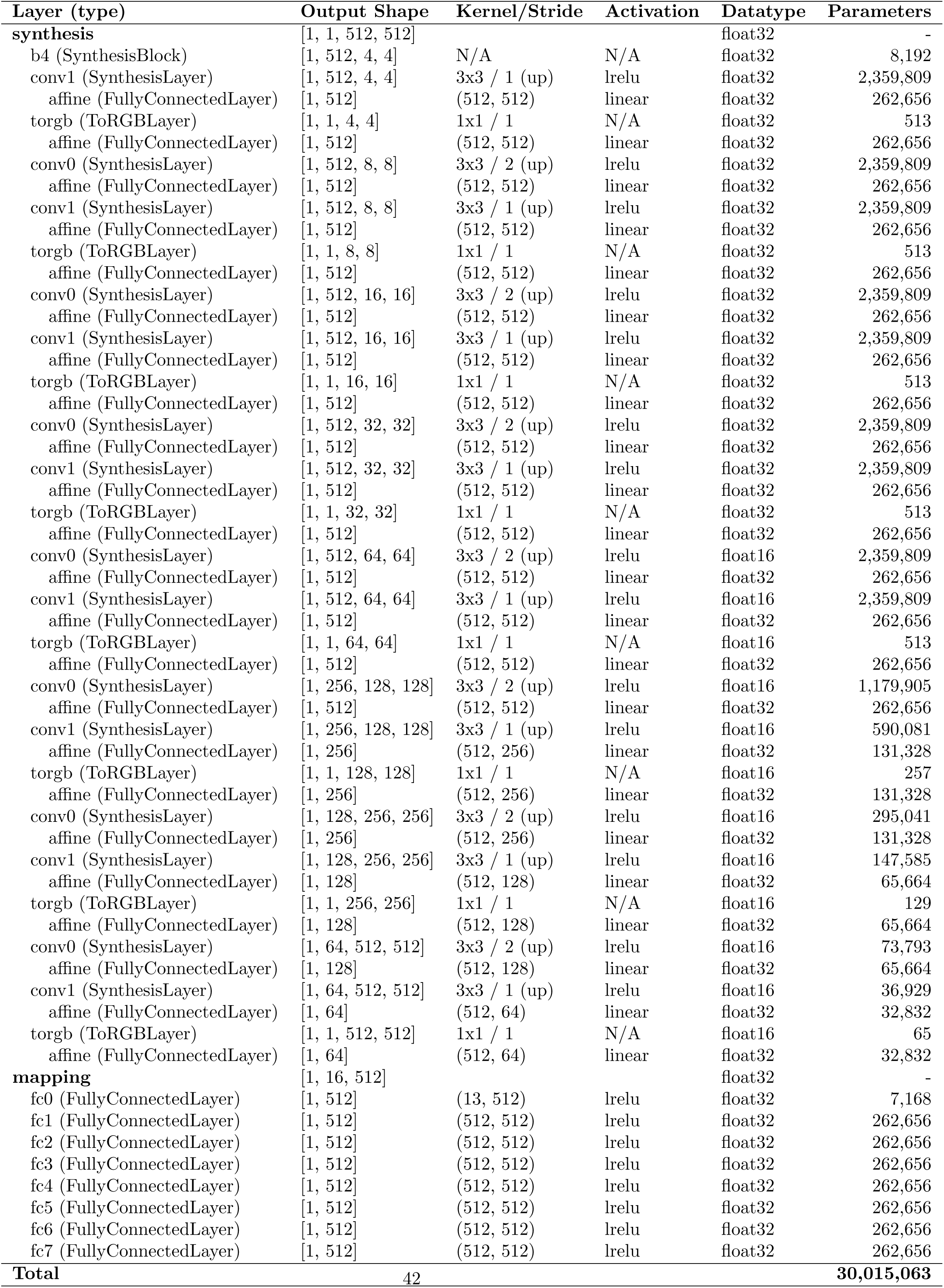
Detailed architecture of the Generator (G) network.

**Table 4:** Detailed architecture of the Discriminator (D) network.

| Layer (type) | Output Shape | Kernel/Stride | Activation | Datatype | Parameters |
| --- | --- | --- | --- | --- | --- |
| fromrgb (Conv2dLayer) | [1, 64, 512, 512] | 1x1 / 1 (up) | lrelu | float16 | 64 |
| conv0 (Conv2dLayer) | [1, 64, 512, 512] | 3x3 / 1 (up) | lrelu | float16 | 36,864 |
| conv1 (Conv2dLayer) | [1, 128, 256, 256] | 3x3 / 2 (down) | lrelu | float16 | 73,728 |
| skip (Conv2dLayer) | [1, 128, 256, 256] | 1x1 / 2 (down) | linear | float16 | 8,192 |
| conv0 (Conv2dLayer) | [1, 128, 256, 256] | 3x3 / 1 (up) | lrelu | float16 | 147,456 |
| conv1 (Conv2dLayer) | [1, 256, 128, 128] | 3x3 / 2 (down) | lrelu | float16 | 294,912 |
| skip (Conv2dLayer) | [1, 256, 128, 128] | 1x1 / 2 (down) | linear | float16 | 32,768 |
| conv0 (Conv2dLayer) | [1, 256, 128, 128] | 3x3 / 1 (up) | lrelu | float16 | 589,824 |
| conv1 (Conv2dLayer) | [1, 512, 64, 64] | 3x3 / 2 (down) | lrelu | float16 | 1,179,648 |
| skip (Conv2dLayer) | [1, 512, 64, 64] | 1x1 / 2 (down) | linear | float16 | 131,072 |
| conv0 (Conv2dLayer) | [1, 512, 64, 64] | 3x3 / 1 (up) | lrelu | float16 | 2,359,296 |
| conv1 (Conv2dLayer) | [1, 512, 32, 32] | 3x3 / 2 (down) | lrelu | float16 | 2,359,296 |
| skip (Conv2dLayer) | [1, 512, 32, 32] | 1x1 / 2 (down) | linear | float16 | 262,144 |
| conv0 (Conv2dLayer) | [1, 512, 32, 32] | 3x3 / 1 (up) | lrelu | float32 | 2,359,296 |
| conv1 (Conv2dLayer) | [1, 512, 16, 16] | 3x3 / 2 (down) | lrelu | float32 | 2,359,296 |
| skip (Conv2dLayer) | [1, 512, 16, 16] | 1x1 / 2 (down) | linear | float32 | 262,144 |
| conv0 (Conv2dLayer) | [1, 512, 16, 16] | 3x3 / 1 (up) | lrelu | float32 | 2,359,296 |
| conv1 (Conv2dLayer) | [1, 512, 8, 8] | 3x3 / 2 (down) | lrelu | float32 | 2,359,296 |
| skip (Conv2dLayer) | [1, 512, 8, 8] | 1x1 / 2 (down) | linear | float32 | 262,144 |
| conv0 (Conv2dLayer) | [1, 512, 8, 8] | 3x3 / 1 (up) | lrelu | float32 | 2,359,296 |
| conv1 (Conv2dLayer) | [1, 512, 4, 4] | 3x3 / 2 (down) | lrelu | float32 | 2,359,296 |
| skip (Conv2dLayer) | [1, 512, 4, 4] | 1x1 / 2 (down) | linear | float32 | 262,144 |
| conv (Conv2dLayer) | [1, 512, 4, 8] | 3x3 / 1 (up) | lrelu | float32 | 2,363,904 |
| fc (FullyConnectedLayer) | [1, 512] | (16384, 512) | lrelu | float32 | 8,389,120 |
| out (FullyConnectedLayer) | [1, 1] | (512, 1) | linear | float32 | 513 |
| **Total** |  |  |  |  | **33,171,009** |

**Table 5:** Pairwise comparisons from Tukey’s HSD test for fidelity scores for expert and non-expert groups. Values are p-values with Hedges’ *g* in parentheses.

| **All** | Real | Ours | StyleGAN2 |
| --- | --- | --- | --- |
| Ours | 0.0000 (0.15) | – | – |
| Stylegan2 | 0.9087 (-0.02) | 0.0002 (0.13) | – |
| Stylegan3 | 0.0000 (1.85) | 0.0000 (2.12) | 0.0000 (1.90) |
| **Non-expert** | Real | Ours | StyleGAN2 |
| Ours | 0.0028 (-0.82) | – | – |
| StyleGAN2 | 0.0000 (2.06) | 0.0000 (1.18) | – |
| StyleGAN3 | 0.0000 (-1.44) | 0.0002 (0.75) | 0.5415 (0.23) |
| **Expert** | Real | Ours | StyleGAN2 |
| Ours | 0.0068 (0.14) | – | – |
| StyleGAN2 | 0.9169 (-0.03) | 0.0481 (0.12) | – |
| StyleGAN3 | 0.0000 (1.62) | 0.0000 (1.86) | 0.0000 (1.68) |

**Table 6:** Pairwise comparisons from Tukey’s HSD test for alignment scores for expert and non-expert groups. Values are p-values with Hedges’ *g* in parentheses.

| **All** | Right Crop | Our | StyleGAN2 |
| --- | --- | --- | --- |
| Ours | 0.0030 (-0.83) | – | – |
| Stylegan2 | 0.0000 (2.04) | 0.0000 (1.21) | – |
| Biological Replicate | 0.0000 (-1.47) | 0.0002 (0.78) | 0.4285 (0.26) |
| **Non-expert** | Right Crop | Ours | StyleGAN2 |
| Ours | 0.0062 (-0.80) | – | – |
| StyleGAN2 | 0.0000 (1.90) | 0.0000 (1.19) | – |
| Biological Replicate | 0.0000 (-1.44) | 0.0003 (0.79) | 0.3469 (0.29) |
| **Expert** | Real Crop | Ours | StyleGAN2 |
| Ours | 0.0015 (0.15) | – | – |
| StyleGAN2 | 0.9918 (-0.01) | 0.0044 (0.14) | – |
| Biological Replicate | 0.0000 (2.11) | 0.0000 (2.41) | 0.0000 (2.13) |

**Table 7:** Pairwise post-hoc test with a Tukey’s HSD test for feature similarity. The numbers provided in the table are p-values, and the numbers in parentheses are effect sizes (Hedge’s *g*).

| **Paired** | Raw | Human | **Random** | Raw | Human |
| --- | --- | --- | --- | --- | --- |
| Human | 0.0092 (0.26) |  | Human | 0.5197 (-0.11) |  |
| Model | 0.0000 (1.16) | 0.0000 (-0.87) | Model | 0.2537 (-0.17) | 0.8752 (0.05) |

## Image and model dataset

DOI 10.6019/S-BIAD2328

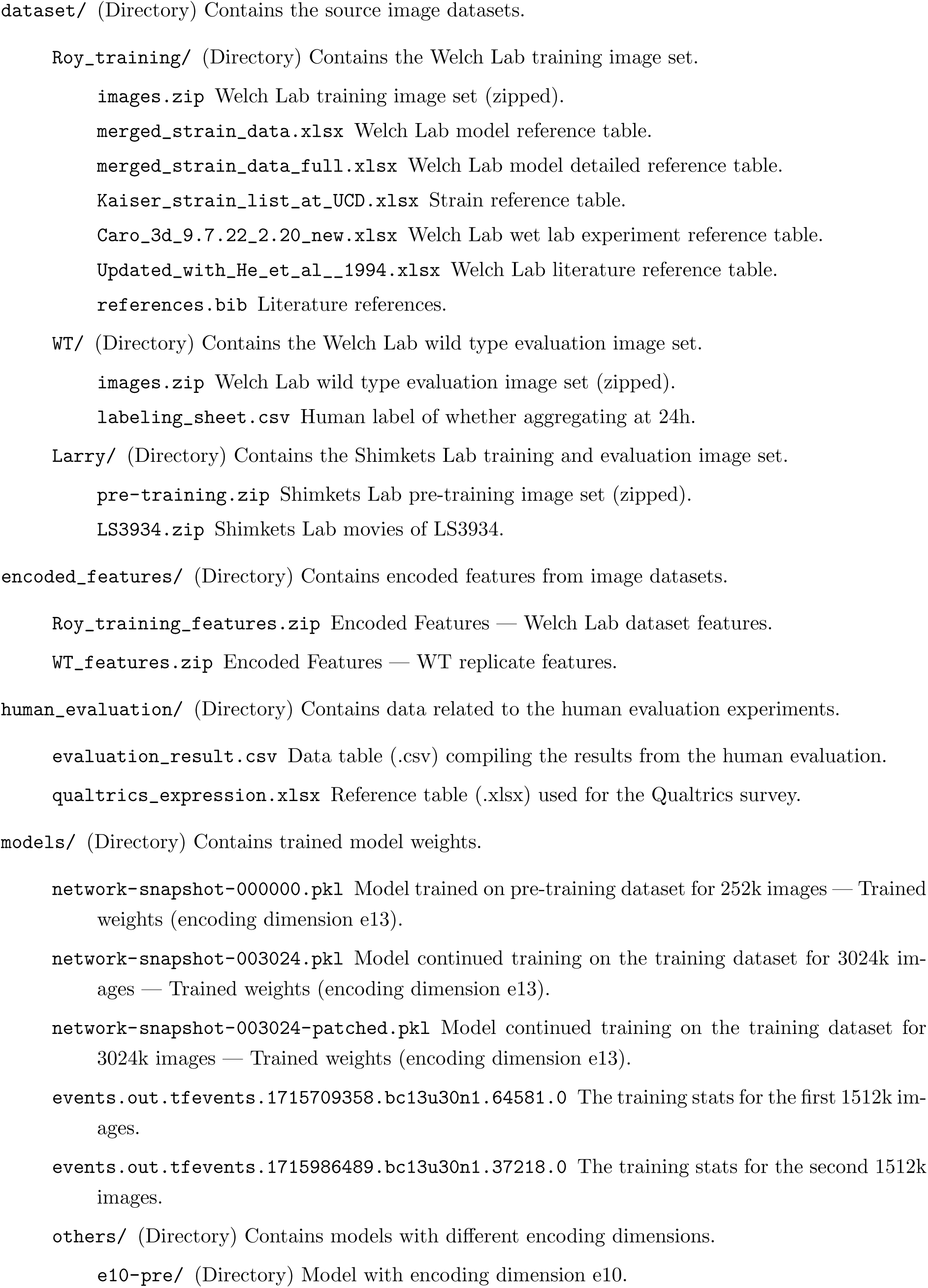
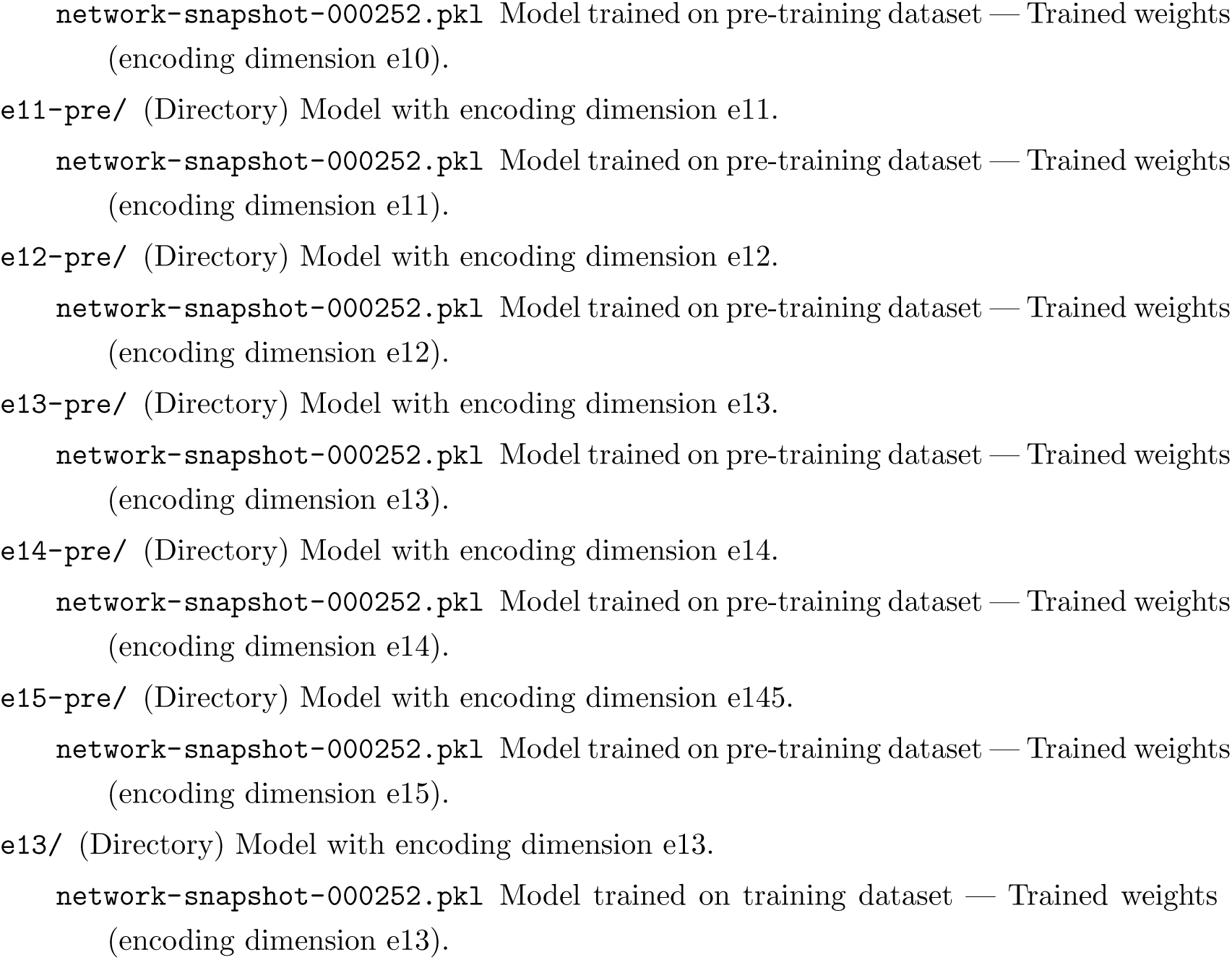

